# Shifted Reverse PAGE: a novel approach based on structure switching for the discovery of riboswitches and aptamers

**DOI:** 10.1101/2022.07.26.501614

**Authors:** Aurélie Devinck, Emilie Boutet, Jonathan Ouellet, Rihab Rouag, Balasubramanian Sellamuthu, Jonathan Perreault

## Abstract

Riboswitches are regulatory sequences composed of an aptamer domain capable of binding a ligand and an expression platform that allows the control of the downstream gene expression based on a conformational change. Current bioinformatic methods for their discovery have various limitations. To circumvent this, we developed an experimental technique to discover new riboswitches called SR-PAGE (Shifted Reverse Polyacrylamide Gel Electrophoresis). A ligand-based regulatory molecule is recognized by exploiting the conformational change of the sequence following binding with the ligand within a native polyacrylamide gel. Known riboswitches were tested with their corresponding ligands to validate our method. SR-PAGE was imbricated within an SELEX to enrich switching RNAs from a TPP riboswitch-based degenerate library to change its binding preference from TPP to thiamine. The SR-PAGE technique allows performing a large screening for riboswitches, search in several organisms and test more than one ligand simultaneously.

## Introduction

Microbial gene expression has to be tightly regulated to adapt to constant changing environments^1^. Cells are able to import molecules from the extracellular environment and their concentration can be crucial to modulate enzymatic pathways^1^. A specific element of the mRNA mostly found in the 5’ untranslated region (UTR) called riboswitch can bind these molecules (ligand) which leads to a change in their secondary structure. This change will affect transcription or translation of the downstream gene. The affected genes often encode for the synthesis or the transportation of the bound molecule^2^. These riboswitches thus comprise an aptamer domain and an expression platform, the structure of which depends on the binding of the aptamer’s cognate ligand. To date, more than 40 riboswitches specifically binding with different metabolites and ligands have been reported^3–9^ in bacteria. Interestingly, the thiamine pyrophosphate (TPP) riboswitches were also observed in archaea, plants and fungi^6^.

The discovery of most riboswitches was made possible by the availability of sequenced genomes for numerous bacteria combined with powerful bioinformatic approaches^10–14^ Potential riboswitches are identified based on the presence of sequence covariation and conservation since the aptamer domain of these regulatory molecules is highly conserved. Covariation occurs when changes in the nucleotide sequence does not affect the RNA secondary structure, reinforcing the importance of its formation^14^ However, bioinformatic tools are limited by sequence availability and their annotations within databases. Also, an alignment of three sequences is theoretically enough to provide covariation that will support a secondary structure prediction and sufficient to warrant experimental evaluation of a putative riboswitch. However, in practice, alignments leading to discovering new riboswitches typically had dozens to hundreds of sequences. The identification of the ligand recognized by the aptamer domain is often facilitated by the nature of the downstream gene. Problematically, gene annotation is sometimes missing or misleading and does not always provide the necessary information to identify the metabolite recognized by the riboswitch. A list of orphan riboswitches for which the associated ligands are currently unknown resulted from bioinformatics-based discovery of numerous putative regulatory RNA structures^10–15^. From the current approach for riboswitch discoveries, it is expected that a very small number of all riboswitch classes have been unveiled^16^. The remaining riboswitches would unfortunately be scarcer, making them much harder to be discovered by bioinformatic tools. Thus, a method which does not rely on sequences available in public databases might be the best way to circumvent this problem.

Some experimental methods have been developed by other groups in recent years to discover new riboswitches. Term-seq enables to screen all the transcription termination within chosen organisms using a high-throughput sequencing approach^17^. This method can be used to identify regulations involving a premature stop of transcription as can be found with some riboswitches. However, this method omits all riboswitches that have an impact on translation. Also, Term-seq cannot differentiate transcription termination caused by a riboswitch from transcription termination caused by a protein, which could be derived from indirect regulation. Another experimental method that can be used to discover new riboswitches is Parallel Analysis of RNA Confirmations Exposed to Ligand Binding (PARCEL)^18^. It is an *in vitro* method to detect structural changes of natural RNA in presence of a ligand by comparing the targeted sites of the RNAse V1, an enzyme that cleaves base-paired regions. A change in the degradation pattern of the RNAse V1 between tested conditions is a good indicator that the ligand induces a change of conformation in the RNA, which is characteristic of riboswitch^18^. However, this technique is limited to regions that are accessible by RNase and only one genome can be analyzed at the time. Another technique derived from SHAPE-MaP^19^ was developed to investigate RNA molecules that interact with ligands. It was recently used to screen hundreds of ligands for their ability to bind RNA molecules^20^. Nevertheless, it could also be used to screen for riboswitches as with PARCEL, but with similar limitations, except perhaps that changes in RNA secondary structure upon binding is resolved to the nucleotide level.

Aptamers for various ligands have successfully been identified based on SELEX (Selective Evolution of Ligands by Exponential Enrichment)^21^, the first ones being RNA aptamers selected against T4 DNA polymerase^22^ and various organic dyes^23^. The general idea is to start with a mix of an extremely diverse library of sequences (~10^16^ sequences) and a target of interest. Unbound sequences are separated from those showing an affinity for the target by a selection method. Selected sequences are amplified for the next round of enrichment. Although the general idea remains the same, many selection methods have been used within a SELEX to isolate aptamers, including capillary electrophoresis (CE-SELEX)^24^, capture-SELEX^25^ and magnetic bead SELEX^26^ to name a few (reviewed in^27,28^). We hypothesized that a SELEX-based approach could also be applied to discover riboswitches. Nonetheless, there are certain restrictions preventing the use of typical SELEX experiments to find natural riboswitches. These regulatory sequences are often engulfing their ligand, guaranteeing that any attempt to immobilize the ligand on a column would cause steric hindrance. Even when riboswitches do not envelop their ligand, the chemical groups chosen for immobilization are likely to be among the groups bound by the potential riboswitch. As for methods that immobilize the library instead of the ligand (e.g., capture-SELEX^25^), it would be difficult to use them to find riboswitches because users need to define a capture sequence within the random region, which is not possible for a library containing unknown natural riboswitches.

We have developed a novel method capable of getting over these obstacles, addressing at the same time the disadvantages of other experimental techniques. This method relies only on the two main characteristics of riboswitches: their ability to bind a ligand and their conformational change upon such binding. These characteristics are evaluated by electrophoresis approach analogous to 2D gels, but instead of running through a second dimension, electrodes are reversed. During the second migration, the RNA runs backwards in presence of a ligand. This allows the detection of a conformational change of the RNA upon binding with a ligand compared to RNA that does not interact with one. We named our technique Shifted Reverse PolyAcrylamide Gel Electrophoresis (SR-PAGE). Our method was validated with known riboswitches by verifying the shifting ability of several constructions. The power of this method was also demonstrated by selecting for a thiamine aptamer from a degenerate library of TPP riboswitches with the SR-PAGE imbricated within a SELEX as a selection method.

## Results

### Validating SR-PAGE with known riboswitches

SR-PAGE can identify riboswitches by taking advantage of the change in the conformation of the RNA sequence following binding with its cognate ligand within a native polyacrylamide gel. Briefly, the method is composed of two native gel migrations: during the first one, the potential regulatory RNA sequences migrate from the wells towards the positive electrode getting separated based on their size and their structure (Fig. 1a). After 24 hours of migration, the gel is unmolded, and the solution of ligand(s) is sprayed on the gel to allow interaction with the RNA molecules (Fig. 1b). The second migration is then started with the same condition as the first migration, but the polarity of the gel is reversed, forcing all RNA sequences to migrate back to the starting point (Fig. 1c). Upon binding with the ligand, the RNA is subjected to a conformational change, either assuming a more compact form that would allow it to migrate pass the well line or creating a more relaxed conformation that would slow down its migration in the gel (Fig. 1c). SR-PAGE can recognize the ligand-based regulatory molecule by selecting those that had a change in migration after the addition of the ligand.

**Fig. 1.**
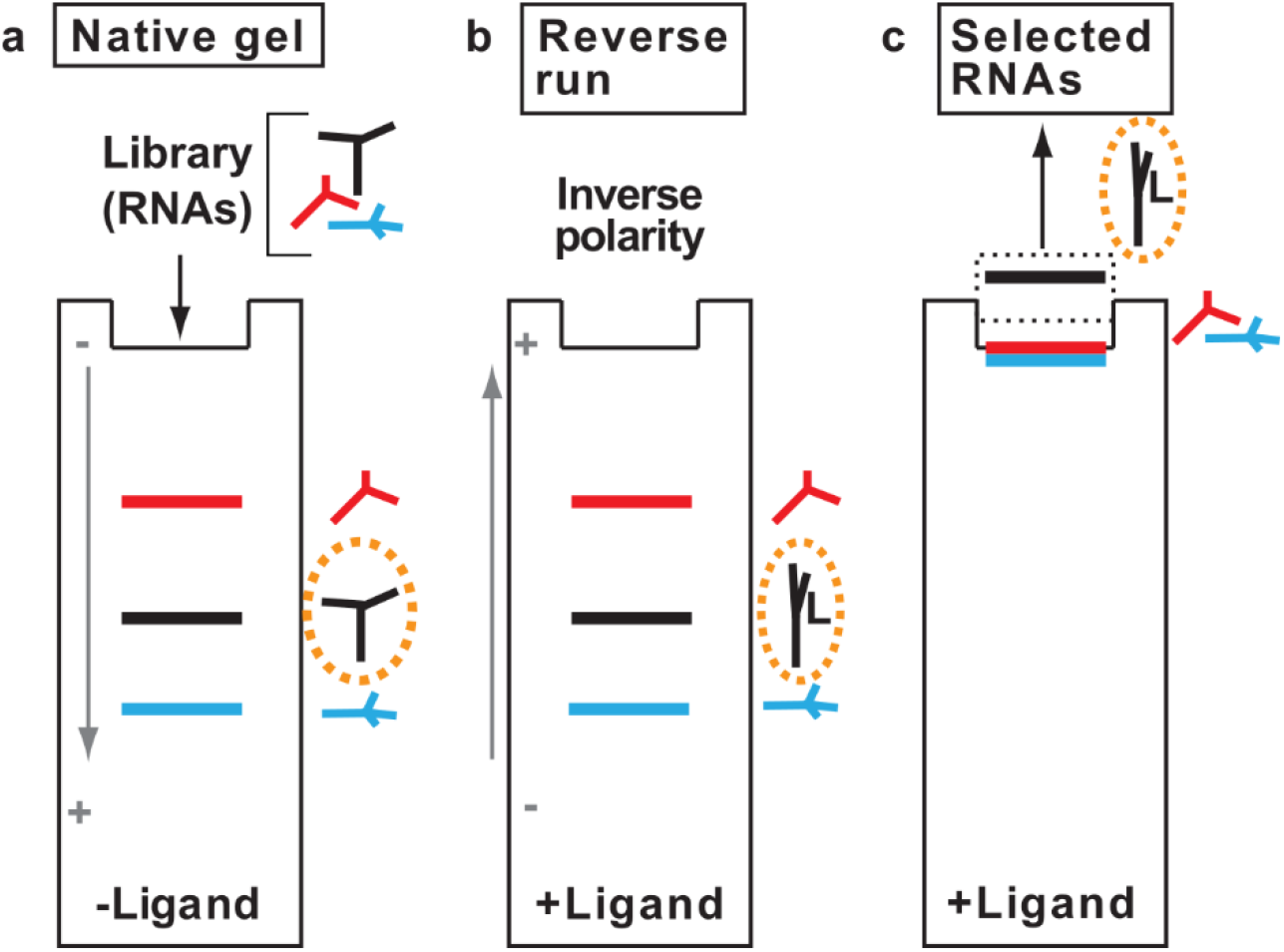
Overview of SR-PAGE. **a**, An RNA library is loaded on a native gel and migrated in absence of ligand. RNAs are separated based on structure and size. **b**, The top glass plate is taken off and the ligand is sprayed on the gel, which only changes the conformation of corresponding riboswitches (within the orange dotted circle) upon binding. The plate is put back on and migration is started with inverted polarity. **c**, The gel is run until the RNA comes back to its starting point. RNAs that interact with the ligand will have a change of migration due to a change in their secondary structure upon ligand binding. A more detailed schematic of the SR-PAGE method is available in the supplementary material (Supplementary Fig. 1).

As a proof of concept, we first wanted to validate that a change in migration upon binding of the ligand could be observed using SR-PAGE with known riboswitches. Up to five different constructions of six known riboswitches were designed, namely the known riboswitches for the following metabolites: flavin mononucleotide (FMN)^29^, fluoride^30^, c-di-GMP (type I)^31^, nickel-cobalt (NiCo)^32^, thiamine pyrophosphate (TPP)^33^ and glycine^34^. The sequence for the glycine riboswitch was based on results by Kwon et al. where a shift was observed in a native gel during migration with glycine^19^ Their constructions varied in length, ranging from only the aptamer to incorporating different portions of the expression platform (Fig. 2 a,b,c; Supplementary Table 1).

**Fig. 2.**
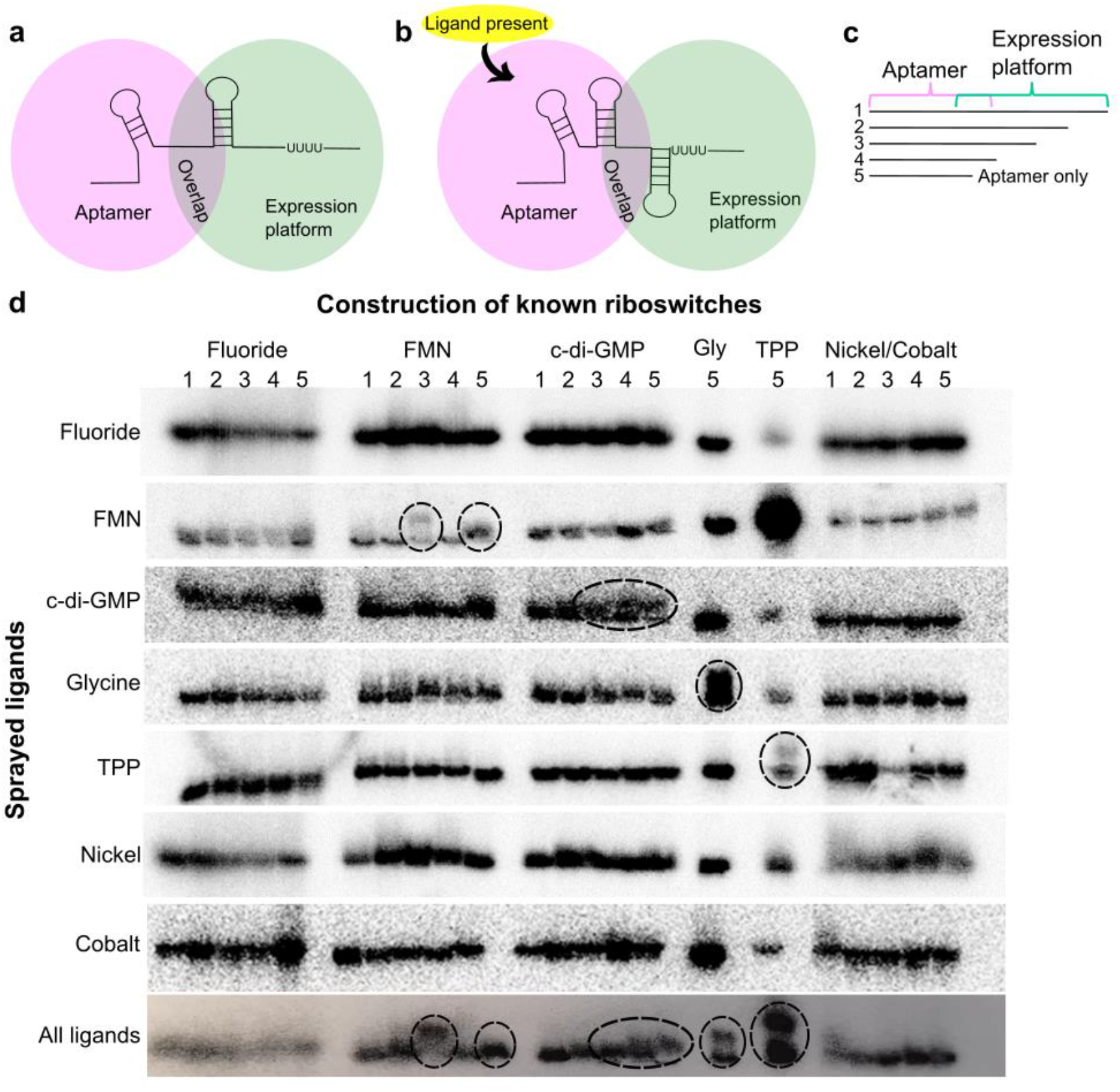
Validation of SR-PAGE with known riboswitches. **a**, Schematic representation of a riboswitch in its unbound conformation, with the aptamer domain and expression platform emphasized in pink and green respectively. The overlap between those two regions is shown. **b**, Schematic of a riboswitch in its bound conformation, where a ligand is bound to the aptamer domain, leading to a change in the secondary and tertiary structure. As an example, the change caused the formation of a Rho-independent transcription terminator in this case. **c**, Representation of how the different constructions (1 to 5) of the known riboswitches were created. **d**, Results from all SR-PAGE. Each group of horizontal bands corresponds to a different gel, where distinct ligands were sprayed into the gel before the second migration (only fluoride, FMN, c-di-GMP, glycine, TPP, nickel, cobalt or all ligands). Five different radiolabeled constructions for each known riboswitch were tested, except for the TPP and glycine riboswitch, where only the aptamer construction was assayed. Observable shifts in migration are circled.

Multiple SR-PAGE experiments were performed, where only one ligand at a time was sprayed at a concentration 100-fold higher than their respective known dissociation constant (*K*_D_) (Fig. 2d). Only the riboswitch constructions corresponding to the sprayed cognate ligands interacted with the metabolites, resulting in a shift of migration (circled in Fig. 2d). RNA molecules that did not interact with the sprayed ligands returned to the starting point in a straight line.

When FMN was sprayed, a significant change in migration for FMN_3 and a slight shift for FMN_5 constructs were observed (Fig. 2d). Small changes in migration for c-di-GMP_3, c-di-GMP_4 and c-di-GMP_5 constructs were also detected when the corresponding ligand was sprayed (Fig. 2d). A change in aptamer migration was also observed for the glycine and TPP aptamers (Fig. 2d) when the corresponding ligands were sprayed. Interestingly, the identical shift in migrations was observed when all ligands were sprayed at the same time on the gel, indicating that SR-PAGE can be used to select for multiple riboswitches with different ligand affinity at the same time (Fig. 2d).

### SR-PAGE to investigate structure changes and expression platforms

We also hypothesized that SR-PAGE could be a means to investigate the expression platform of a riboswitch. It could be used to select for the construction that results in a bigger change in migration, implying it has a bigger conformational change. FMN_3 and FMN_5 were further analyzed to understand why different shift levels were observed between constructions of the same riboswitch: they technically both have similar ability to interact with the metabolite, since they are constituted of the same aptamer domain (Fig. 3). The Mfold web service^35^ was employed to analyze the free energy of both constructions in their bound and unbound forms. To mimic the bound conformation, sequences were forced to adopt the FMN aptamer domain structure^36^ using constraints (constraints used to force the formation of the aptamer can be found in Supplementary material, Table 2). The equilibrium constant was then measured from the difference in free energies between the two conformations.

**Fig. 3.**
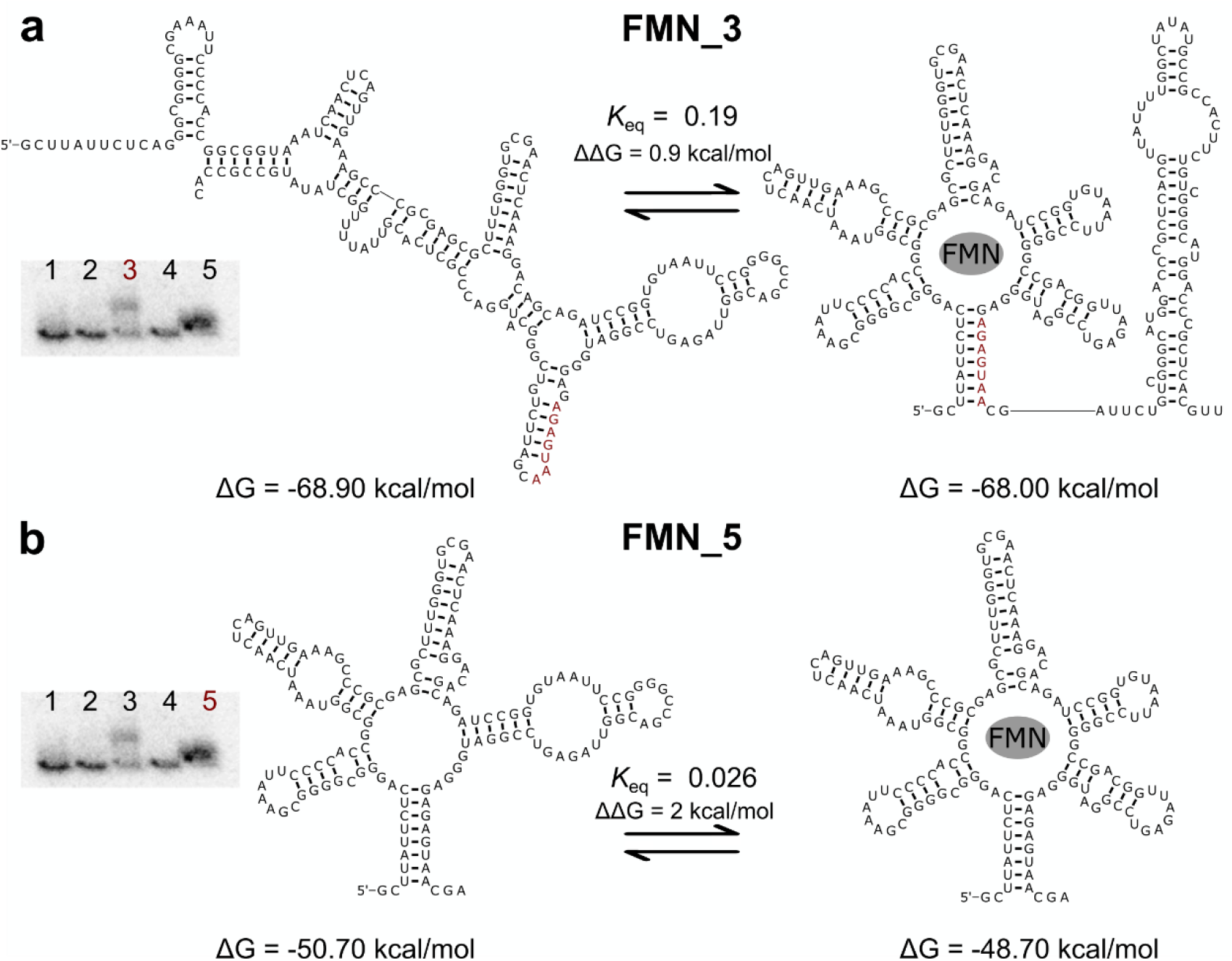
Shifting constructions 3 and 5 of the FMN riboswitch have similar free energies for their bound and unbound states. **a,b** Predicted structures, free energies (ΔG) and equilibrium constant (*K*_eq_) of the unbound RNA and ligand-bound state of FMN_3 and FMN_5, respectively. Nucleotides emphasized in red can basepair to form the basal stem of the aptamer, leading to a mutually exclusive conformation. Shifts observed by SR-PAGE are shown, with corresponding construction number highlighted in red for a and b.

For FMN_3, the RNA is only slightly more stable in its unbound conformation (unconstrained) compared to when constraints are used to force the aptamer conformation, with free energies of −68.9 kcal/mol and −68 kcal/mol respectively (Fig. 3a). Our interpretation is that during the first migration, the FMN_3 assumes its more stable unbound conformation (Fig. 3a, left) which can potentially interconvert with the aptamer-containing structure (Fig. 3a, right), with a preference of 5:1 for the former compared to the latter, according to the calculated equilibrium constant. However, when the FMN ligand is sprayed unto the gel, not only does the equilibrium change, but the “aptamer state” of the RNA (capable of interacting with the ligand) also likely has changes in its tertiary structure. The change in structure from the more stable one (on the left) to the aptamer state results in the shift of migration observed with SR-PAGE (Fig. 3a). Even if the so-called unbound state should be favored, since the free energy of both structures is similar, the presence of the ligand suffices to permit the switch. The same situation seems to apply for FMN_5 (Fig. 3b), with free energies of −50.70 kcal/mol and −48.70 kcal/mol for the unbound and bound conformations, respectively. However, the two structures appear closely related, which may explain the less prominent shift. Previous work also demonstrated that the aptamer of the FMN riboswitch is pre-formed in the absence of the metabolite at physiological concentrations of magnesium^37^. In this case, in spite of a *K*eq suggesting a ratio of ~40:1 of the most stable vs. the aptamer state, the closeness of both structures may permit an easier switch in equilibrium, but also a more complete one, as suggested by the quasi-absence of a non-shifted band for FMN_5.

Both FMN_3 and FMN_5 structures are more stable in their unbound state, so the equilibrium constant favors in each case this conformation (in absence of ligand), represented by a *K*_eq_ value smaller than 1. Some of the RNA molecules were still able to adopt the ligand-binding conformation, since the *K*_eq_ value is close to 1. A similar analysis of the free energies and equilibrium constants of ligand-bound state compared to free RNA for all constructions of riboswitches FMN, fluoride, c-di-GMP and nickel-cobalt is available in the supplementary material (Supplementary Figs. 2–5). The larger differences in free energies observed for all the constructions of fluoride and nickel-cobalt riboswitches may explain that none of the tested constructs shifted (Supplementary Figs. 2 and 5). Moreover, the equilibrium constant for most of these cases had very small values, meaning that the “aptamer state” is likely almost absent. As for c-di-GMP_3-4-5, FMN_3 and FMN_5, most free energies are very similar with *K*_eq_ closer to 1, which explains that the constructs all seem to shift, but the shifts are more subtle given the similarity between the predicted bound and unbound states.

### Selection of thiamine aptamer from degenerate libraries of TPP riboswitch

As a proof of concept that the SR-PAGE can be used as a method of selection in a SELEX, the aptamer sequence from the TPP riboswitch was partially randomized to create three libraries possessing each 4,096, 1,048,576 and 68,719,476,736 possible sequence combinations (Fig. a). Libraries 1, 2 and 3 contained 6, 10 and 18 degenerated nucleotides respectively. Briefly, RNA libraries were run on native polyacrylamide gel electrophoresis during SR-PAGE. Thiamine was added to the second run (reversed polarity) of SR-PAGE, first at 2 mM (generations 1 to 4) and in later rounds of selection, at a lower concentration of 20 μM (generation 5) and 200 nM (generations 7 to 10) to increase the stringency. Shifted RNAs were purified and amplified, then loaded on gel together with TPP (as a negative selection ligand) for the next round of selection, where thiamine is again added only for the reverse migration. We therefore selected for RNA molecules that preferentially bind thiamine rather than its native TPP. After four, five, seven and ten rounds of selection, selected sequences were cloned and also analyzed by Illumina sequencing^38^ to identify the enriched sequences. RNAs were selected based on their shifting abilities during the SR-PAGE (Fig. 4b).

**Fig. 4.**
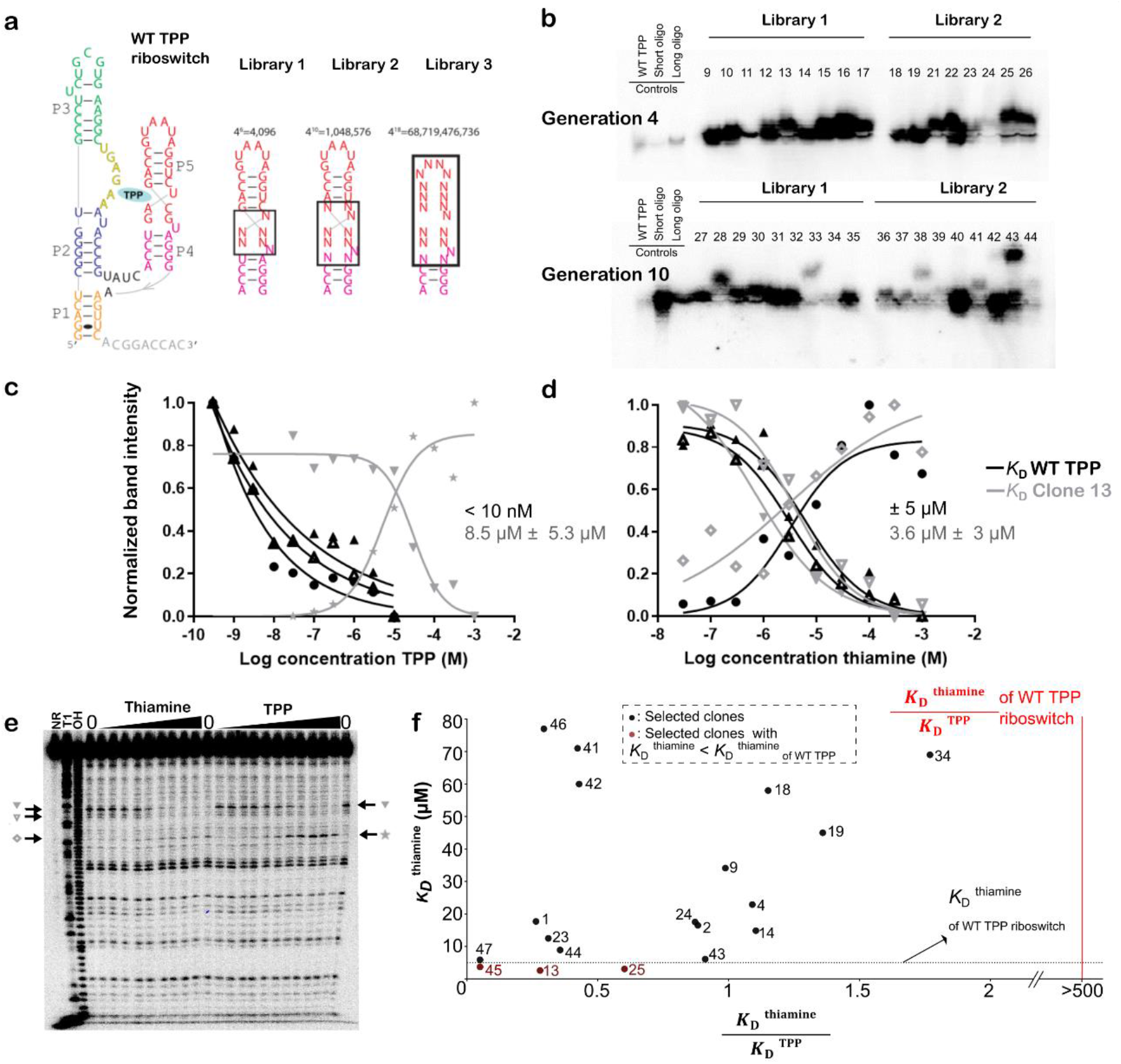
Selection of a thiamine aptamer from degenerated libraries of the TPP riboswitch using the SR-PAGE as a selection tool within a SELEX. **a**, Degenerate libraries of the TPP riboswitch were created. Libraries 1, 2 and 3 had 6, 10 and 18 degenerated nucleotides respectively within stem P5. **b**, Shifting abilities as a result of SR-PAGE from libraries 1 and 2 with clones selected during generations 4 and 10 of the SELEX. TPP was present during the first migration within the running buffer (10 μM), and thiamine (20 μM) was sprayed on the gel before the second migration. RNA molecules can be observed since they are radioactively labeled. As controls, a short and long oligos as well as the WT TPP riboswitch were migrated on the same gel. **c**, *K*_D_ curve for WT TPP riboswitch (in black) and selected sequence 13 from library 1 after four generations of SELEX (in gray) with TPP. **d**, *K*_D_ curve for WT TPP riboswitch (in black) and selected sequence 13 from library 1 after four generations of SELEX (in gray) with thiamine. **e**, Inline probing gel of clone 13. The arrows represent the bands used for *K*_D_ calculation (graphed in c). Tested concentrations for thiamine and TPP ranged from 30 nM to 1 mM. **f**, Affinity of all selected sequences with thiamine and TPP. All dots represent individual selected sequences, where those emphasized in red are clones with a better affinity for thiamine than the WT TPP riboswitch. The dotted horizontal line depicts the *K*_D_ of the WT TPP riboswitch for thiamine. The vertical red line represents the ratio of *K*_D_ for thiamine over *K*_D_ for TPP of the WT TPP riboswitch.

A smaller dissociation constant (*K*_D_) represents a higher affinity of a molecule to its ligand. While the wildtype aptamer has a *K*_D_ smaller than 10 nM for TPP and of 4 μM ± 1.9 μM for thiamine in our tested conditions (Fig. c), selected sequences using SR-PAGE as a selection method kept their binding affinity for thiamine and lost affinity for TPP through rounds of SELEX (Fig. 4f). *K*_D_ of candidates were determined by in-line probing^14^. One of the best clones (clone 13) showed a slightly improved binding for thiamine molecules, with a *K*_D_ of 3 μM ± 1.3 μM for thiamine and 7.8 μM ± 3.5 μM for TPP (Fig. 4c, d, e). Other clones (1, 2, 9, 13, 23, 24, 25, 41, 42, 43, 44, 45, 46 and 47) also showed that the aptamer recognize thiamine with a better affinity than TPP molecules (Fig. 4f and Supplementary Table 3). Only clones where a *K*_D_ could be estimated from in-line probing are shown in the figure. Sequences of all selected clones as well as their *K*_D_ with thiamine and TPP when measurable can be found in Supplementary Table 3 (nomenclature is explained in Supplementary Fig. 6).

In library 3, the P5 stem of the WT TPP riboswitch was initially replaced by random nucleotides, therefore only a small proportion of sequences are expected to form a similar stem, and none are expected to have the WT internal loop, since we do not select for TPP binding (Fig. 4a). After four and seven rounds of SELEX, selected libraries were analyzed by Illumina sequencing. Two of the clones (45 and 47) that we tested from this initial library show no binding to the TPP molecule up to a concentration of 1 mM (Fig. 4f). In spite of the fact that we completely degenerated stem P4-P5 (except for two bp), we apparently still selected for a stem (Supplementary Fig. 7a). Overall, secondary structure prediction for enriched sequences (i.e., more than three reads/sequence) showed that we selected for the formation of a stem in P5, as opposed to random sequences that do not typically form a stem (Supplementary Fig. 7; and Supplementary Table 4). Because the thiamine binding is not greatly improved (as suggested by *K*_D_s similar to that of WT), this rather suggests that the stem was selected because it improved the shift between unbound states and thiamine-bound. Indeed, angles between stems are known to critically impact how RNA runs in a native PAGE^39^.

One concern of using the SR-PAGE within a SELEX was the impact of the adapters necessary for the PCR amplification of the selected sequences in a SELEX round. Our results show that the adapters can indeed prevent riboswitch shifting using our positive controls (Supplementary Fig. 8). The addition of oligonucleotides complementary to the adapters helps to overcome this problem (Supplementary Fig. 8). All sequences for the aptamers and oligonucleotides can be found in the supplementary materials (Supplementary Table 1).

## Discussion

In this paper, we presented the Shifted Reverse PAGE (SR-PAGE), a novel method to study riboswitches and aptamers going beyond the limits of bioinformatics and other existing experimental methods. This technique was validated with several known riboswitches, namely c-di-GMP-I, FMN, TPP and glycine, including when multiple ligands were used at the same time. We also demonstrated that the SR-PAGE could be imbricated within an SELEX as a method of selection, allowing us to identify degenerated TPP riboswitch sequences that lost their affinity for TPP and improved their affinity for thiamine, as confirmed by in-line probing.

A shift in migration for all known riboswitches was not always detectable, like for the fluoride and nickel-cobalt riboswitches for example (Fig. 2d). This can be explained by the tested constructions. Indeed, as demonstrated with the FMN_3 and FMN_5 constructions (Fig. 3), a few nucleotides make the difference between an RNA that shifts and an RNA that does not shift, likely due to a subtle equilibrium of structures between the conditions with and without ligands. Thus, a slight difference in sequence can result in a big change in terms of secondary structure. Even if all tested constructions technically contained the aptamer domain, it does not necessarily mean that the aptamer was forming. We hypothesized that a shift was not observed for all known riboswitches simply because the RNA sequence resulting in a bigger change of secondary structure upon ligand binding was not tested, or likely because the difference in free energy between the unbound and bound structure was too large to be overcome in other cases. The difference in predicted free energies was in average of 1.78 kcal/mol in constructions where a shift in migration was observed (FMN_3, FMN_5, c-di-GMP_3-4-5; Fig. 2d), whereas it was on average 12.1 kcal/mol for those where no change in migration was seen (FMN_1-2-4, c-di-GMP_1-2, NiCo_1-2-3-4-5, fluoride_1-2-3-4-5; Fig. 2d). In these latter cases, the free energy difference between the unbound state and the ligand-bound state is too large to be overcome by the ligand. The equilibrium constant values in cases where no shift was observed were also small (typically <10^-3^), suggesting that the secondary structures probably could not adopt the ligand-bound conformation and/or that the energy of ligand-binding was insufficient to overcome the gap between ΔGs. A shift was observed in cases where the equilibrium constant was closer to 1, suggesting the RNA structure could overcome the difference in free energy to be able to bind the ligand. This is the induced-fit model^40^, where the addition of the ligand favors a change in secondary structure of a given RNA molecule. On another end, the addition of the ligand could favor RNA molecules that are already in the proper secondary structure, which is the conformational selection model^41^. In this case, the dynamic interchange of structures within the gel would be locked into the “aptamer state” upon addition of ligand to explain the change in migration. It is likely that combinations of both models occur in many cases, including also more subtle changes of the tertiary structure which could nevertheless lead to detectable shifts in the gel.

The amplitude of the shift (the height of the shift) appears to correlate with the level of structural difference between the unbound and the bound structure, where similar secondary structure result in a smaller change in migration. This was observed for example in FMN_5, where the shift was less pronounced than that of FMN_3, since it was already in a secondary structure closely related to that of the aptamer (Fig. 3), as well as c-di-GMP constructions, which were similar for the most part. Therefore, one of the limitations of SR-PAGE is that we might miss riboswitches that are able to bind a ligand, but the conformational change is not drastic enough to result in a detectable change in migration. However, this limitation can be used to our advantage, since the SR-PAGE can be used to identify constructions that will produce larger shifts and gain insight on the expression platform, further improving our knowledge of existent riboswitches and aptamers, as suggested with the analysis of the FMN riboswitch construction (Fig. 3).

We have also used SR-PAGE within a SELEX strategy to select riboswitches with a modified affinity for their ligands. The SR-PAGE selected RNA molecules based on two properties: first, for their ability to preferentially bind thiamine rather than TPP molecules, and for a change in the RNA conformation involving a shift by SR-PAGE upon that binding. More than forty sequences identified from that SELEX were tested separately by in-line probing. Out of all these sequences, 14 showed a better affinity for thiamine compared to that of the TPP molecule (*K*_D_ thiamine / *K*_D_ TPP <1; compared to>500 for WT). Moreover, three clones (13, 25 and 45) had a better affinity for thiamine than the WT TPP riboswitch (5 μM). This experiment successfully demonstrated that SR-PAGE could be used as a selection step within a SELEX to select for affinity modified and/or enhanced aptamers.

Now that the SR-PAGE method has been validated, it could be employed to identify potential novel riboswitches starting with genomic libraries of intergenic sequences of bacteria, for example. The validation of the SR-PAGE technique presents a new approach to discover riboswitches, which has advantages over the normally used bioinformatics methods. Only intergenic sequences are studied with bioinformatics analysis, whereas SR-PAGE allows researchers to look at the entire genomic sequences. Less complex RNA secondary structure can be overlooked when using bioinformatics, which SR-PAGE would not disregard. For example, mini-ykkC (aka guanidine-II) was deemed too simple to be a riboswitch until it was assayed with guanidine^42^. Moreover, novel regulatory structures will be directly linked to their ligand, contrary to bioinformatics, which often leads to orphan riboswitches, where the specificity of the ligand is not known^11,12,14,15,43^. The SR-PAGE selects for sequences that change conformation upon binding of a ligand. Therefore, potential riboswitches still need be validated *in-vivo* using for example riboswitch-reporter fusion assay to confirm it has an impact on the downstream gene expression. Nevertheless, the use of SR-PAGE is not intended to overshadow bioinformatics techniques, but rather to complement them, since both have their advantages. This technique can be applied to several types of ligands such as coenzymes, amino acids and metal ions. Each new discovery will provide new insights into the biochemical capacity of RNA and open opportunities for technological advances.

## Methods

### PCR construction of riboswitches

Forward and reverse primers were designed to amplify the different constructions of the FMN and the fluoride riboswitches from the genomes of *Escherichia coli (E. coli)* and *Burkholderia thailandensis (B. thailandensis)* respectively, with the addition of the T7 promoter. For the glycine, TPP, nickel-cobalt and c-di-GMP class I riboswitches, PCR assembly was used to create the template using Primerize^44^. The different constructions of the c-di-GMP riboswitch were then created in the same manner as for the FMN and the fluoride riboswitches. All oligonucleotides used are listed in Supplementary Table 1. The PCR reactions were done with 2.5 U Hot Start Taq Polymerase (Qiagen/203203) with Hot Start buffer (1X), dNTPs (200 μM), forward and reverse primers (1 μM) completed with Milli-Q water. For assembly PCR, the end primers have a concentration of 1 μM and the internal oligonucleotides have a concentration of 0.1 μM.

Three libraries derived from *E. coli thiM* TPP riboswitch were designed to obtain an aptamer that binds thiamine much better than TPP. To do so, the pyrophosphate binding pocket within P5 was randomized to different degrees with degenerated primers (Fig. 4a) to create the libraries 1, 2 and 3. These oligonucleotides are also listed in the Supplementary Table 1.

### *In vitro* RNA transcription

For SR-PAGE, all PCR products of the known riboswitches as well as those from the clones derived from the SELEX with degenerated TPP riboswitches were radiolabeled with [α-^32^P] UTP (Perkin Elmer) during *in vitro* transcription using T7 RNA polymerase for 3 hours at 37 °C (20 μL PCR products [~400 ng], 20 μL 5X transcription buffer [400 mM HEPES-KOH, pH 7.5, 120 mM MgCl_2_, 10 mM spermidine, 200 mM DTT], 0.001 U pyrophosphatase (Roche), 0.5 μL RNAse inhibitor Ribolock (Thermo Fischer/EO0382) [40 U/ μL], T7 polymerase [100 U], 5 mM ATP, 5 mM GTP, 5 mM CTP, 1 mM UTP and 5 μCi [α-^32^P] UTP). The volume was completed to 100 μL with Milli-Q water. The transcribed RNA was precipitated at −80°C with 0.1 volume of sodium acetate 3 M (pH 5.2) and 2 volumes of 100% chilled ethanol for at least 2 hours. The precipitated RNA was resuspended in RNAse-free Milli-Q water and purified on a 6% denaturing polyacrylamide gel electrophoresis (8 M urea PAGE). Samples were loaded on the gel with 2X denaturing loading buffer (0.05% bromophenol blue, 0.05% xylene cyanol, 10 mM EDTA [pH 8], 95% formamide). The expected size bands corresponding to our RNA were eluted in the elution buffer (0.3 M NaCl) and the eluate was precipitated as described before. Radiolabeled RNAs were resuspended in 250 μL of Milli-Q sterilized water.

For in-line probing, RNA molecules were radiolabeled at their 5’ end. *In vitro* transcription was performed as before, but with equal concentrations of all rNTPs, since radioactive [α-^32^P] UTP were not used. After an *in vitro* transcription, 5’ ends were dephosphorylated using 5 U of Antarctic phosphatase (New England BioLabs/M0289S). A mix containing the RNA sample (~10 pmol), Antarctic phosphatase buffer 1X and 0.5 μL RNase inhibitor Ribolock (Thermo Fischer/EO0382). The volume was adjusted to 20 μL with sterilized Milli-Q water. The reaction was incubated at 37 °C for 20 minutes and deactivated at 65°C for 5 minutes. RNA molecules were radiolabeled at their 5’ end in a reaction with ~10 pmol dephosphorylated RNA samples, 0.2 μL T4 Polynucleotide Kinase (PNK) enzyme (New England BioLabs/M0201S), 1 μL 10X T4 PNK buffer and 2 μL [γ-^32^P] ATP (Perkin Elmer). The volume was adjusted to 10 μL with water and incubated at 37°C for 1 hour. Labeled RNA samples were purified on a 6% denaturing PAGE, imaged and purified as described before.

### SR-PAGE gel preparation

A slab gel system, typical of sequencing gels, was used. To mold the gel, we used glass plates of dimensions (38 cm x 45 cm) for the larger one and of (38 x 43 cm) for the smaller one. The smaller glass plate was treated with a solution of Rain-X ®, a hydrophobic solution, whereas the larger glass plate was treated with a solution of potassium chloride (KOH) and methanol (5 g of KOH in 100 mL of methanol), a hydrophilic solution. These steps become important later when it is time to disassemble the gel to ensure that the gel sticks to the larger hydrophilic plate rather than on the smaller hydrophobic one. The plates are then assembled leaving a 2 cm gap at the bottom of the gel between the small and large plate (Supplementary Fig. 1a). Spacers are placed between the two plates, representing the thickness of the gel (approximately 0.8 mm). A 250 mL solution of native polyacrylamide gel (29:1) 10% was prepared with a final TBMg concentration of 1X (0.09 M Tris base, 0.09 M boric acid and 5 mM Mg(CH_3_COO)_2_ pH 8.0 at ambient temperature). The volume was completed with sterilized Milli-Q water. A volume of 50 mL of this solution was kept at 4°C for later use to maintain the exact same percentage of acrylamide for the solution used to cover the void left by the well removal. To polymerize the gel, 2 mL of 10% ammonium persulfate (APS, Bio-Rad) and 70 μL of tetramethylethylenediamine (TEMED, Bioshop) was added to the remaining 200 mL preparation of the native polyacrylamide gel. Once the gel was poured, the gel was left to polymerize for at least 30 minutes. The assembly of the SR-PAGE was made as shown in Supplementary Fig.1b. To allow a good circulation of the buffer throughout the gel, a peristaltic pump system was installed to carry the buffer from the bottom tank to top tanks. The buffer was also allowed to flow from the top tank to the bottom one via a tube (Supplementary Fig. 1b). Pre-migration was carried out overnight at 450 V with TBMg 1X as running buffer. From this point on, all the steps were carried in a 4°C room.

RNA samples were prepared in TBMg 1X, 3,3 μL native blue 6X loading dye (40% sucrose, 0.05 % bromophenol blue, 0.05% xylene cyanol). Different controls were used to monitor proper migration. We used radioactive RNAs of different sizes (short and long) to show that regardless of the size, the RNA comes back to the starting point if it did not interact with the ligand. Another control is a Cy3 fluorescent oligonucleotide to visualize the migration within the gel with the naked eye (Supplementary Table 1). Finally, depending on the SR-PAGE assay being run, a riboswitch construct known to shift in native gels was often used as a positive control (validated by our preliminary results). These controls were prepared in the same way as the RNA samples. The wells were cleaned with a syringe to remove all potential bubbles and unpolymerized acrylamide in the wells before loading the samples and controls. RNA molecules were separated according to their size and structure for 24 hours in the 10% native polyacrylamide gel at 450 V.

### SR-PAGE reverse migration

After the first migration of 24 hours, the glass plates were carefully separated. A 50 mL solution containing the ligands of interest was prepared. The final vaporized concentration of each known riboswitch’s ligand is equivalent to hundred times their respective known *K*_D_. The sprayed concentrations for each ligand were as follows: 6 mM FH_4_KO_2_^45^, 5 μM FMN sodium salt, hydrate^46^ (Cayman chemical company), 3 μM c-di-GMP sodium salt^47^ (Sigma-Aldrich), 10 mM glycine^48^ (Bioshop), 3 μM TPP chloride^33^ (Sigma-Aldrich), 3 mM NiCl_2_^32^ (Fischer), 3 mM CoCl_2_^32^ (Fisher) in TBMg 1X. When we worked with metal ions, 10 mM L-glutathione (C_10_H_17_N_3_O_6_S) (Sigma-Aldrich) was also added to the sprayed solution as a reducing agent. The volume was adjusted to 50 mL with sterilized Milli-Q water. This solution was then sprayed directly onto the gel and left in contact for ten minutes (Supplementary Fig.1c). It is important to spray the solution across all the surface of the gel as uniformly as possible. The wells were cut out and the space left by the removal of these wells was carefully dried with Kimwipes® paper (Kimtech). TEMED solution (70 μL) was added to the base of the wells to help seal the junction between the old and newly added acrylamide solution. The glass plates were reassembled, this time perfectly aligning the bottom of the two plates. The previously prepared native polyacrylamide solution was degassed and used to fill the space left by the wells, taking care to add the ligands sprayed into the solution at a final concentration 10 times lower than that present in the sprayed solution, since we do not rely on passive diffusion in the gel in this case. To allow the gel to polymerize, 700 μL of 10% APS and 70 μL of TEMED were added to the 50 mL native polyacrylamide gel preparation. Note that these steps were carried at 4°C, so higher concentrations of APS and TEMED are required for the gel to polymerize. The gel was left to polymerize for at least 30 minutes. The SR-PAGE was then reassembled as before, but the polarity of the gel was reversed, with the negative electrode now at the bottom and the positive at the top. The migration was restarted at 450 V (Supplementary Fig.1c). Using a marker, the previous well line was delineated on the glass plates to use as a reference point to know when to stop the second migration.

Migration was stopped once the fluorescent oligonucleotide reached the wells. Due to a decreased resistance, the fluorescent oligonucleotide reached the baseline in a little less time than the first migration. This time could range anywhere between 19h to 23h, so it was important to monitor closely the SR-PAGE when we reached the end of the second migration. When it was not necessary to purify RNA from the gel, as for radioactively labeled riboswitches from Figs. 2d and 4b, the top of the gel was cut off and then dried using a gel dryer (model 583 Biorad, with a vacuum pump CVC3000 Vacuubrand). The gel was then exposed overnight on a phosphor screen and visualized with a Typhoon FLA9500 (GE Healthcare Life Sciences). For SELEX assays, gel strips were cut. Band 0 corresponds to the well baseline, band 1 corresponds to ~2 mm above the well up to 1 cm above the wells and band 2 corresponds to all the rest of the top of the gel; then the RNAs were eluted as described before.

### SELEX of thiamine-binding TPP-derived riboswitches

When SR-PAGE was imbricated in an SELEX, like for the selection of a modified aptamer starting from degenerate libraries of TPP riboswitches, gel regions corresponding to “shifted-RNA” were cut out. RNAs were eluted in elution buffer overnight and ethanol-precipitated like described before. Reverse transcription (RT) was performed with M-MuLV reverse transcriptase (NEB) according to manufacturer instructions with a short oligonucleotide to prevent annealing of the oligonucleotides on RNAs that lost a few nucleotides at the 3’ end during the migration (and would thus run faster in the reverse run, co-migrating with “shifted” RNA). Half of the RT was used as a PCR template for a 100 μL reaction with conditions like library generation, but with 30 cycles for the first generation, and then progressively less for subsequent generations. RNAs were transcribed *in vitro* to start a new round of selection with an SR-PAGE like described before.

During selection, 10 μM of TPP was added to the running buffer of the SR-PAGE. For the second migration, thiamine was sprayed on the gel, with a decreasing concentration as the SELEX generation increased to improve stringency. Therefore, 2 mM of thiamine was sprayed on the gel for generation 1 through 4. Then, 20 μM of thiamine was added for generation 5. Finally, 200 nM of thiamine was sprayed for the last cycles (generations 7 to 10).

### In-line probing and dissociation constant determination

Sequences from the initial library or from generation 4, 5, 7 and 10 were either cloned with the pGEM®-T kit (Promega) to assay individual sequences and/or sequenced by Illumina (Centre d’expertises et de services Genome Québec). After analysis of Illumina sequencing results, sequences that were enriched during the cycles were selected. Individual sequences were amplified by PCR, transcribed to RNA by *in vitro* transcription, purified on 6% 8 M urea PAGE, eluted and precipitated as described before.

In-line probing reactions were performed with different concentrations of ligands ranging from 30 nM to 1 mM in a final volume of 20 μL containing 20 mM MgCl_2_, 50 mM Tris pH 8.3 (at 25°C) and 100 mM KCl at room temperature for approximately 40 hours. For the WT TPP riboswitch, the concentrations of ligands were ranging from 30 nM to 1 mM for thiamine and from 1 nM to 30 μM for TPP. The volume of the reaction was adjusted with Milli-Q water. The resulting reaction was quenched with 2X loading buffer. Three ladders (NR: No Reaction, i.e., RNA in water; T1: RNAse T1 which cleaves at guanine bases; and OH: alkaline reaction which cleaves all positions) were made to map the RNA sequence. For the T1 ladder, 10 μL of the radioactively labeled RNA was incubated in 50 mM of sodium citrate, 3 μL of formamide 100% and 4 μL of Milli-Q water. The solution was incubated at 56°C for 2 minutes before 1.5 μL of T1 RNAse 1 U/μL was added to the T1 solution. It was left to react at 56°C for 5 minutes. For the alkaline digestion (OH), 10 μL of the radioactively labeled RNA was incubated with 8 μL of Milli-Q water and 50 mM sodium carbonate at 90°C for 90 seconds. The non-reacted ladder contained only 10 μL of the radioactively labeled RNA with Milli-Q water to a final volume of 20 μL. All reactions for the creation of the ladders were stopped with 20 μL of 2X loading buffer. Samples and ladders were loaded on an 8% urea-PAGE for approximately 3 hours at 50 W^49^. The gel was then dried and exposed on a phosphor screen overnight before being visualized on a Typhoon FLA9500 (GE Healthcare Life Sciences).

For dissociation constant determination, the intensity of the lanes and modulating bands were quantified using the ImageQuant TL software (GE Healthcare Life Sciences). These bands were normalized to give values between 0 and 1 according to minimum and maximum intensity of the bands. The *K*_D_ representation and determination were done with logarithmic curves on GraphPad 7 software^42^.

### Free energy calculation and RNA secondary structure

Free energy of all RNA sequences were assessed using the website service of Mfold accessible at www.unafold.org^35^. RNAs were forced in the aptamer state using constraint based on consensus of secondary structure in Rfam^50^. All constraints applied to Mfold are available in supplementary material (Supplementary Table 2). RNA secondary structures were designed using the Stockholm file generated by Mfold^35^ as input for R2R^51^.

### Equilibrium constant calculation

The equilibrium constants between the bound and unbound conformation of the tested RNA molecules were calculated using the following formula:

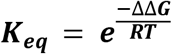

The ΔΔG (in joules) is the difference in free energy (ΔG) of the bound and unbound conformation. The gas constant (R) is 8.314 J/mol·K. The temperature is 277 K, since all SR-PAGE experiments have been carried out at 4 ° C.

## Supporting information

Supplementary Material_SR-PAGE

## Acknowledgements

Authors wish to thank Gaël Montagne for technical help. Early work on SR-PAGE was started within the Breaker lab, JP wishes to thank Ronald Breaker for this, as well as members of the Breaker lab, including Rüdiger Welz who had performed gel shifts with the glycine riboswitch, providing the first positive control for SR-PAGE.

## Author Contributions

A.D. and E.B. contributed equally to this work. J.P. conceived the SR-PAGE method. A.D., E.B., J.O., R.R., B.S. and J.P. optimized the method. J.O. realized the SELEX of thiamine-binding TPP-derived riboswitches. A.D. and E.B. performed all other experiments presented in the article. Figures were created by J.P., J.O., A.D. and E.B. Manuscript was written by J.P., A.D. and E.B. Project was supervised by J.P. Manuscript was revised by all authors.

## Supplementary information

is available for this paper.

## Correspondence and request for materials

should be addressed to J.P.

## References

1 Hennecke, H. Regulation of bacterial gene expression by metal–protein complexes. Mol. Microbiol. 4, 1621–1628 (1990).

2 Serganov, A. & Nudler, E. A decade of riboswitches. Cell 152, 17–24 (2013).

3 Barrick, J. E. et al. New RNA motifs suggest an expanded scope for riboswitches in bacterial genetic control. Proceedings of the National Academy of Sciences 101, 6421–6426 (2004).

4 Mandal, M., Boese, B., Barrick, J. E., Winkler, W. C. & Breaker, R. R. Riboswitches control fundamental biochemical pathways in Bacillus subtilis and other bacteria. Cell 113, 577–586 (2003).

5 Mandal, M. et al. A glycine-dependent riboswitch that uses cooperative binding to control gene expression. Science 306, 275–279 (2004).

6 McCown, P. J., Corbino, K. A., Stav, S., Sherlock, M. E. & Breaker, R. R. Riboswitch diversity and distribution. Rna 23, 995–1011 (2017).

7 Ramesh, A. & Winkler, W. C. Magnesium-sensing riboswitches in bacteria. RNA biology 7, 77–83 (2010).

8 Sherlock, M. E., Sudarsan, N. & Breaker, R. R. Riboswitches for the alarmone ppGpp expand the collection of RNA-based signaling systems. Proceedings of the National Academy of Sciences 115, 6052–6057 (2018).

9 Winkler, W. C., Nahvi, A., Sudarsan, N., Barrick, J. E. & Breaker, R. R. An mRNA structure that controls gene expression by binding S-adenosylmethionine. Nature Structural & Molecular Biology 10, 701–707 (2003).

10 Stav, S. et al. Genome-wide discovery of structured noncoding RNAs in bacteria. BMC microbiology 19, 1–18 (2019).

11 Weinberg, Z. et al. Identification of 22 candidate structured RNAs in bacteria using the CMfinder comparative genomics pipeline. Nucleic acids research 35, 4809–4819 (2007).

12 Weinberg, Z. et al. Detection of 224 candidate structured RNAs by comparative analysis of specific subsets of intergenic regions. Nucleic acids research 45, 10811–10823 (2017).

13 Weinberg, Z., Nelson, J. W., Lünse, C. E., Sherlock, M. E. & Breaker, R. R. Bioinformatic analysis of riboswitch structures uncovers variant classes with altered ligand specificity. Proceedings of the National Academy of Sciences 114, E2077–E2085 (2017).

14 Weinberg, Z. et al. Comparative genomics reveals 104 candidate structured RNAs from bacteria, archaea, and their metagenomes. Genome biology 11, 1–17 (2010).

15 Greenlee, E. B. et al. Challenges of ligand identification for the second wave of orphan riboswitch candidates. RNA Biol 15, 377–390 (2018).

16 Breaker, R. R. Prospects for riboswitch discovery and analysis. Molecular cell 43, 867–879 (2011).

17 Dar, D. et al. Term-seq reveals abundant ribo-regulation of antibiotics resistance in bacteria. Science 352, aad9822 (2016).

18 Tapsin, S. et al. Genome-wide identification of natural RNA aptamers in prokaryotes and eukaryotes. Nature communications 9, 1–10 (2018).

19 Siegfried, N. A., Busan, S., Rice, G. M., Nelson, J. A. & Weeks, K. M. RNA motif discovery by SHAPE and mutational profiling (SHAPE-MaP). Nature methods 11, 959–965 (2014).

20 Zeller, M. J. et al. SHAPE-enabled fragment-based ligand discovery for RNA. Proceedings of the National Academy of Sciences 119, e2122660119 (2022).

21 Irvine, D., Tuerk, C. & Gold, L. Selexion: Systematic evolution of ligands by exponential enrichment with integrated optimization by non-linear analysis. Journal of molecular biology 222, 739–761 (1991).

22 Tuerk, C. & Gold, L. Systematic evolution of ligands by exponential enrichment: RNA ligands to bacteriophage T4 DNA polymerase. science 249, 505–510 (1990).

23 Ellington, A. D. & Szostak, J. W. In vitro selection of RNA molecules that bind specific ligands. nature 346, 818–822 (1990).

24 Zhu, C., Yang, G., Ghulam, M., Li, L. & Qu, F. Evolution of multi-functional capillary electrophoresis for high-efficiency selection of aptamers. Biotechnology advances 37, 107432 (2019).

25 Boussebayle, A., Groher, F. & Suess, B. RNA-based capture-SELEX for the selection of small molecule-binding aptamers. Methods 161, 10–15 (2019).

26 Espelund, M., Stacy, R. & Jakobsen, K. A simple method for generating single-stranded DNA probes labeled to high activities. Nucleic acids research 18, 6157 (1990).

27 Bayat, P. et al. SELEX methods on the road to protein targeting with nucleic acid aptamers. Biochimie 154, 132–155 (2018).

28 Darmostuk, M., Rimpelova, S., Gbelcova, H. & Ruml, T. Current approaches in SELEX: An update to aptamer selection technology. Biotechnology Advances 33, 1141–1161, doi:https://doi.org/10.1016/j.biotechadv.2015.02.008 (2015).

29 Pedrolli, D. et al. The ribB FMN riboswitch from Escherichia coli operates at the transcriptional and translational level and regulates riboflavin biosynthesis. The FEBS journal 282, 3230–3242 (2015).

30 Breaker, R. New insight on the response of bacteria to fluoride. Caries research 46, 78–81 (2012).

31 Sudarsan, N. et al. Riboswitches in eubacteria sense the second messenger cyclic di-GMP. Science 321, 411–413 (2008).

32 Furukawa, K. et al. Bacterial riboswitches cooperatively bind Ni2+ or Co2+ ions and control expression of heavy metal transporters. Molecular cell 57, 1088–1098 (2015).

33 Rentmeister, A., Mayer, G., Kuhn, N. & Famulok, M. Conformational changes in the expression domain of the Escherichia coli thiM riboswitch. Nucleic acids research 35, 3713–3722 (2007).

34 Kwon, M. & Strobel, S. A. Chemical basis of glycine riboswitch cooperativity. Rna 14, 25–34 (2008).

35 Zuker, M. Mfold web server for nucleic acid folding and hybridization prediction. Nucleic acids research 31, 3406–3415 (2003).

36 Serganov, A., Huang, L. & Patel, D. J. Coenzyme recognition and gene regulation by a flavin mononucleotide riboswitch. nature 458, 233–237 (2009).

37 Vicens, Q., Mondragón, E. & Batey, R. T. Molecular sensing by the aptamer domain of the FMN riboswitch: a general model for ligand binding by conformational selection. Nucleic acids research 39, 8586–8598 (2011).

38 Bentley, D. R. et al. Accurate whole human genome sequencing using reversible terminator chemistry. nature 456, 53–59 (2008).

39 Lafontaine, D. A., Norman, D. G. & Lilley, D. M. Structure, folding and activity of the VS ribozyme: importance of the 2-3-6 helical junction. The EMBO Journal 20, 1415–1424 (2001).

40 Heppell, B. et al. Molecular insights into the ligand-controlled organization of the SAM-I riboswitch. Nature chemical biology 7, 384–392 (2011).

41 Haller, A., Rieder, U., Aigner, M., Blanchard, S. C. & Micura, R. Conformational capture of the SAM-II riboswitch. Nature chemical biology 7, 393–400 (2011).

42 Sherlock, M. E., Malkowski, S. N. & Breaker, R. R. Biochemical validation of a second guanidine riboswitch class in bacteria. Biochemistry 56, 352–358 (2017).

43 Meyer, M. M. et al. Challenges of ligand identification for riboswitch candidates. RNA biology 8, 5–10 (2011).

44 Tian, S. & Das, R. Primerize-2D: automated primer design for RNA multidimensional chemical mapping. Bioinformatics 33, 1405–1406 (2017).

45 Baker, J. L. et al. Widespread genetic switches and toxicity resistance proteins for fluoride. Science 335, 233–235 (2012).

46 Howe, J. A. et al. Atomic resolution mechanistic studies of ribocil: A highly selective unnatural ligand mimic of the E. coli FMN riboswitch. RNA biology 13, 946–954 (2016).

47 Smith, K. D. et al. Structural basis of ligand binding by a c-di-GMP riboswitch. Nature structural & molecular biology 16, 1218 (2009).

48 Huang, L., Serganov, A. & Patel, D. J. Structural insights into ligand recognition by a sensing domain of the cooperative glycine riboswitch. Molecular cell 40, 774–786 (2010).

49 Regulski, E. E. & Breaker, R. R. in post-transcriptional gene regulation 53–67 (Springer, 2008).

50 Kalvari, I. et al. Rfam 14: expanded coverage of metagenomic, viral and microRNA families. Nucleic Acids Research 49, D192–D200 (2021).

51 Weinberg, Z. & Breaker, R. R. R2R-software to speed the depiction of aesthetic consensus RNA secondary structures. BMC bioinformatics 12, 1–9 (2011).

