## Supplementary Material_SR-PAGE for "Shifted Reverse PAGE: a novel approach based on structure switching for the discovery of riboswitches and aptamers"

#### Supplementary figures

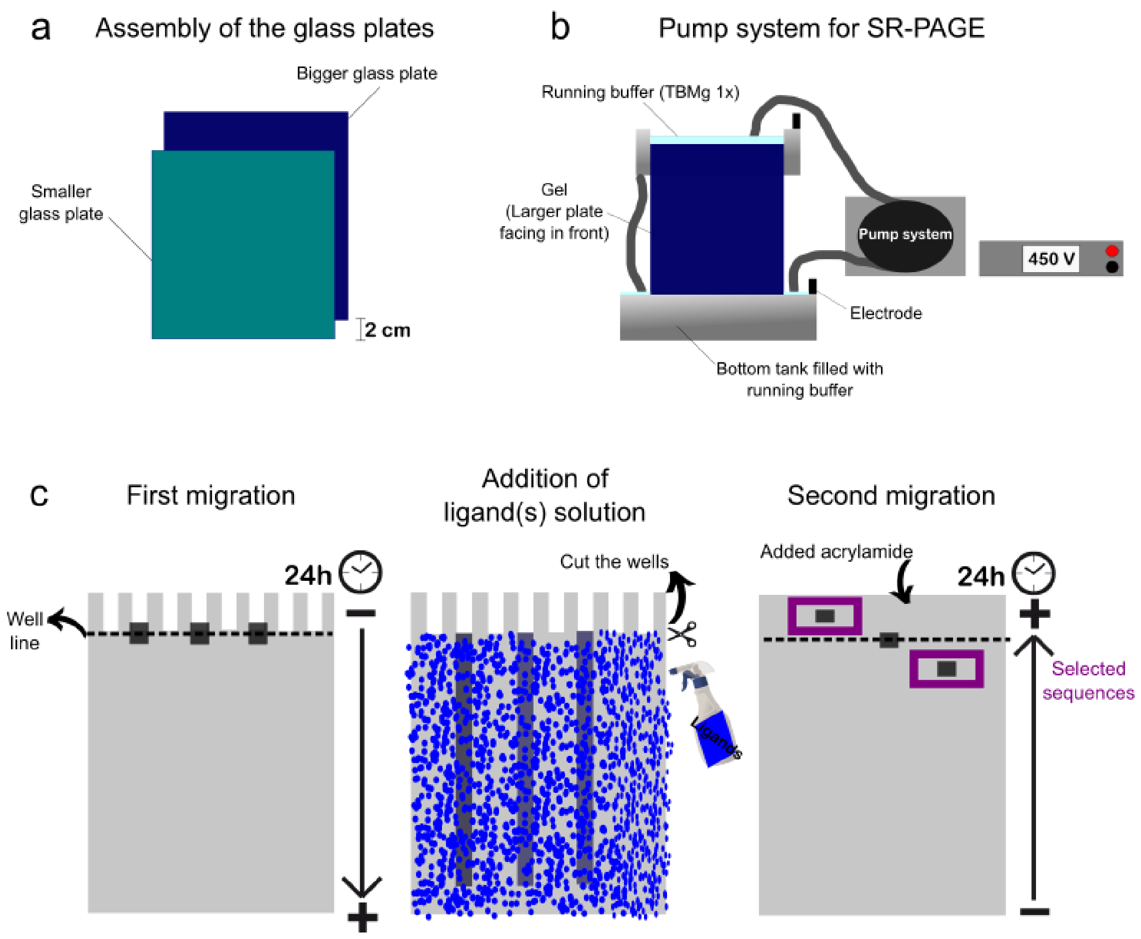

**Supplementary Fig. 1| SR-PAGE method.**

**a,** When the gel is poured between the two glass plates, the smaller glass plate is placed 2 cm below the bigger glass plates as shown. **b**, Schematic representation of a SR-PAGE assembly with the pump system. **c,** First migration of the RNA sequences within a native polyacrylamide gel for 24 hours. Secondly, the gel is small plate is removed and the ligand solution is sprayed unto the gel. The wells are cut. The space left by the removal of the wells is filled by the leftover native polyacrylamide so that the reverse run could cover above the wells. Finally, the second migration of the RNA sequence where the polarity of the electrode is inverted. The sequences that had a change in migration (that did not end at the well line after the second migration) are selected.

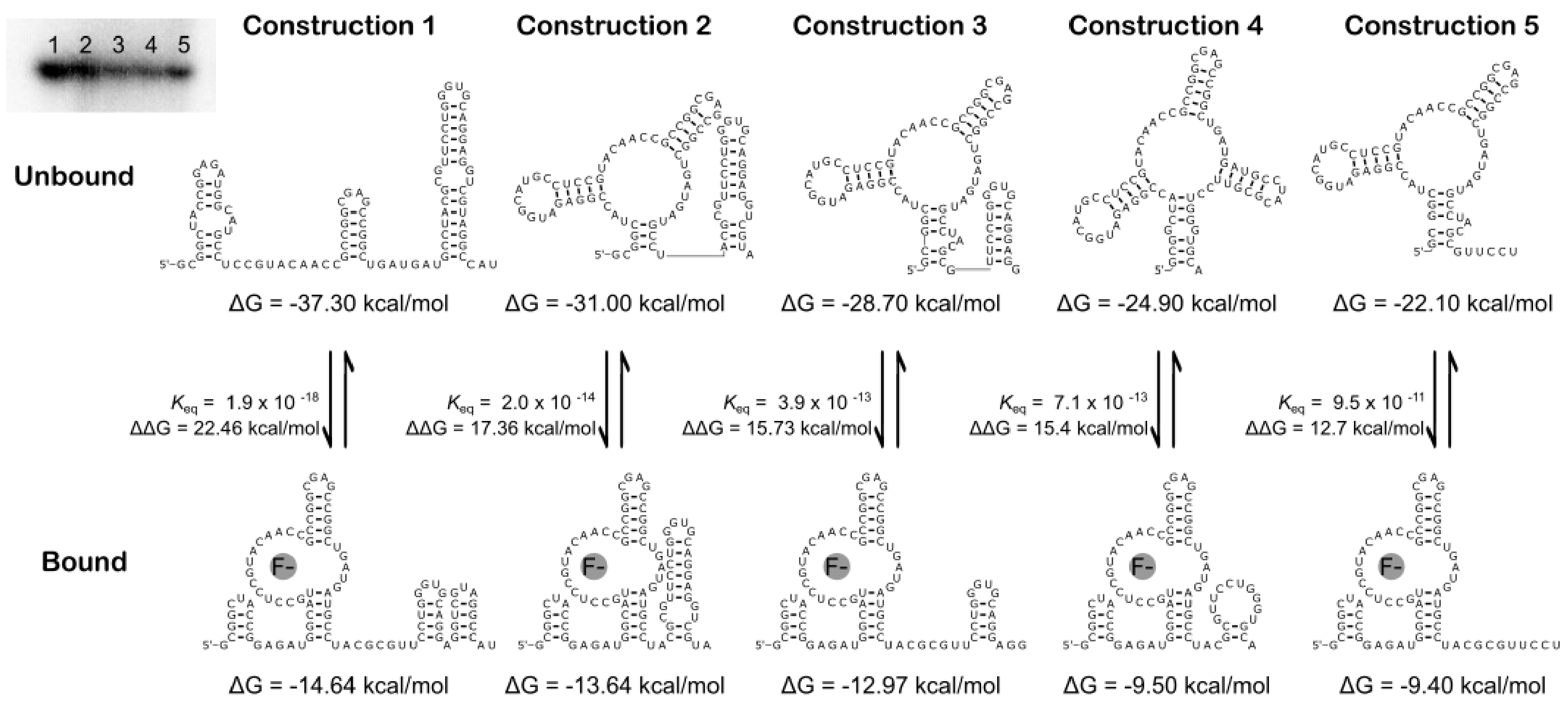

**Supplementary Fig. 2 | Free energy of all different constructions of the fluoride riboswitch used in the SR-PAGE experiment in their bound (constrained) and unbound (unconstrained) conformations.**

For Supplementary figures 2-5, the structures were obtained with Mfold using constraints described in Supplementary Table 3 and corresponding to the known folding of the aptamer bound-conformation according to available atomic resolution data of the given riboswitch aptamer domain (as the structure of the expression platform was not solved for these examples), within the boundaries of Mfold’s capacity (i.e. precluding non-canonical base pairs and other similar tertiary interactions). The unbound version simply corresponds to the unconstrained Minimum Free Energy (MFE) structure predicted by Mfold. Secondary structures were generated with R2R.

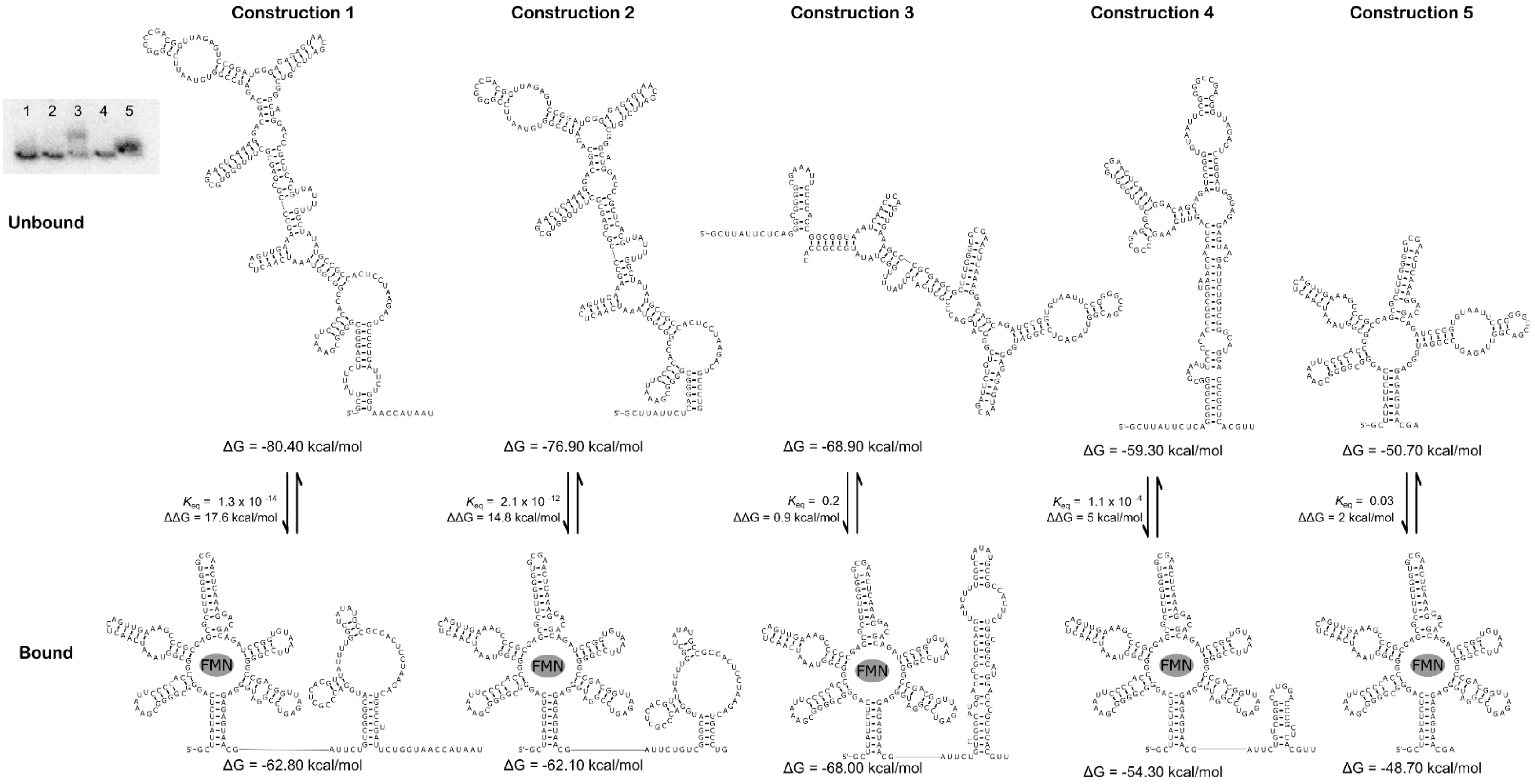

**Supplementary Fig. 3 | Free energy of all different constructions of the FMN riboswitch used in the SR-PAGE experiment in their bound (constrained) and unbound (unconstrained) conformations.**

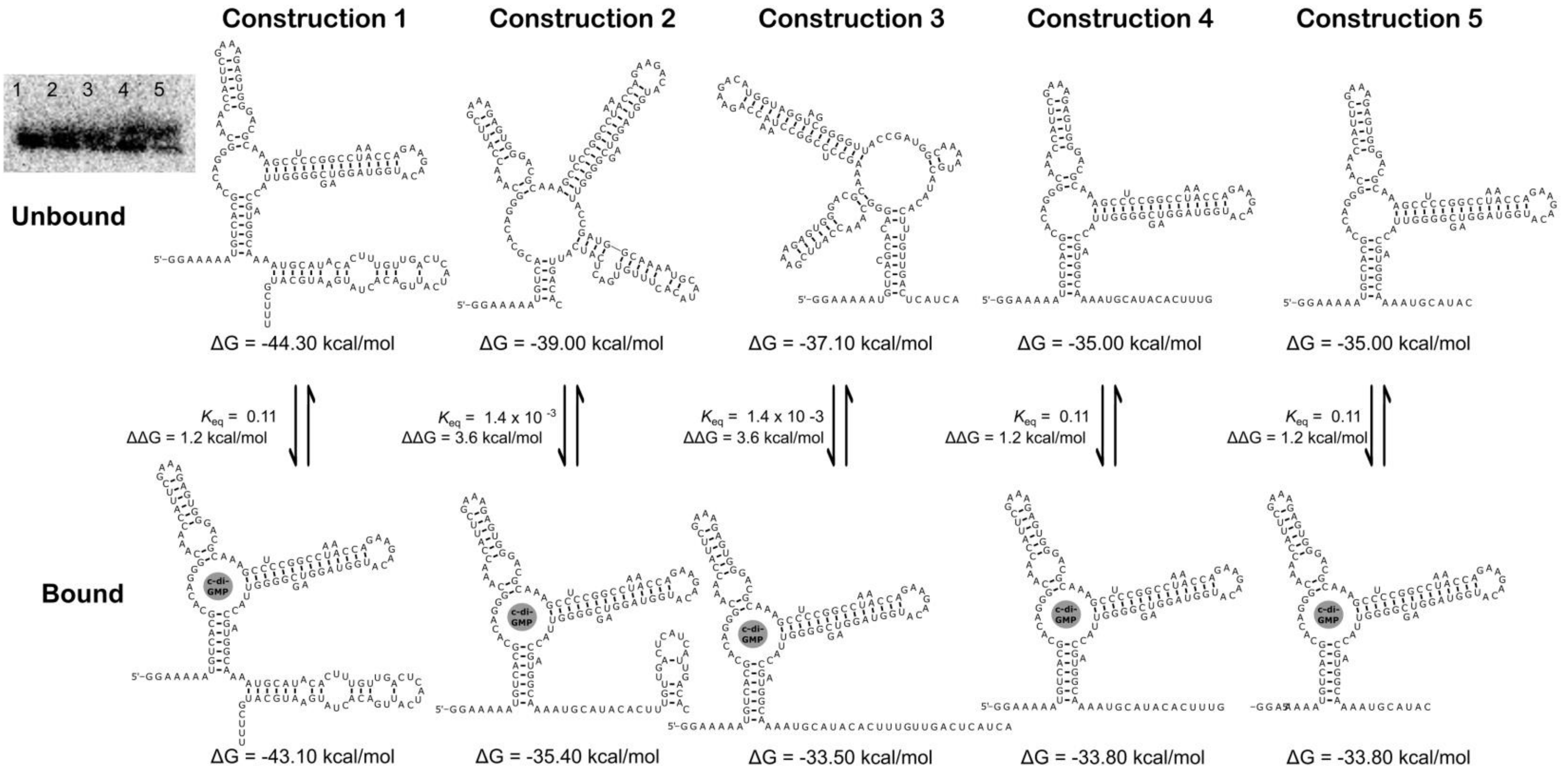

**Supplementary Fig. 4 | Free energy of all different constructions of the c-di-GMP I riboswitch used in the SR-PAGE experiment in their bound (constrained) and unbound (unconstrained) conformations.**

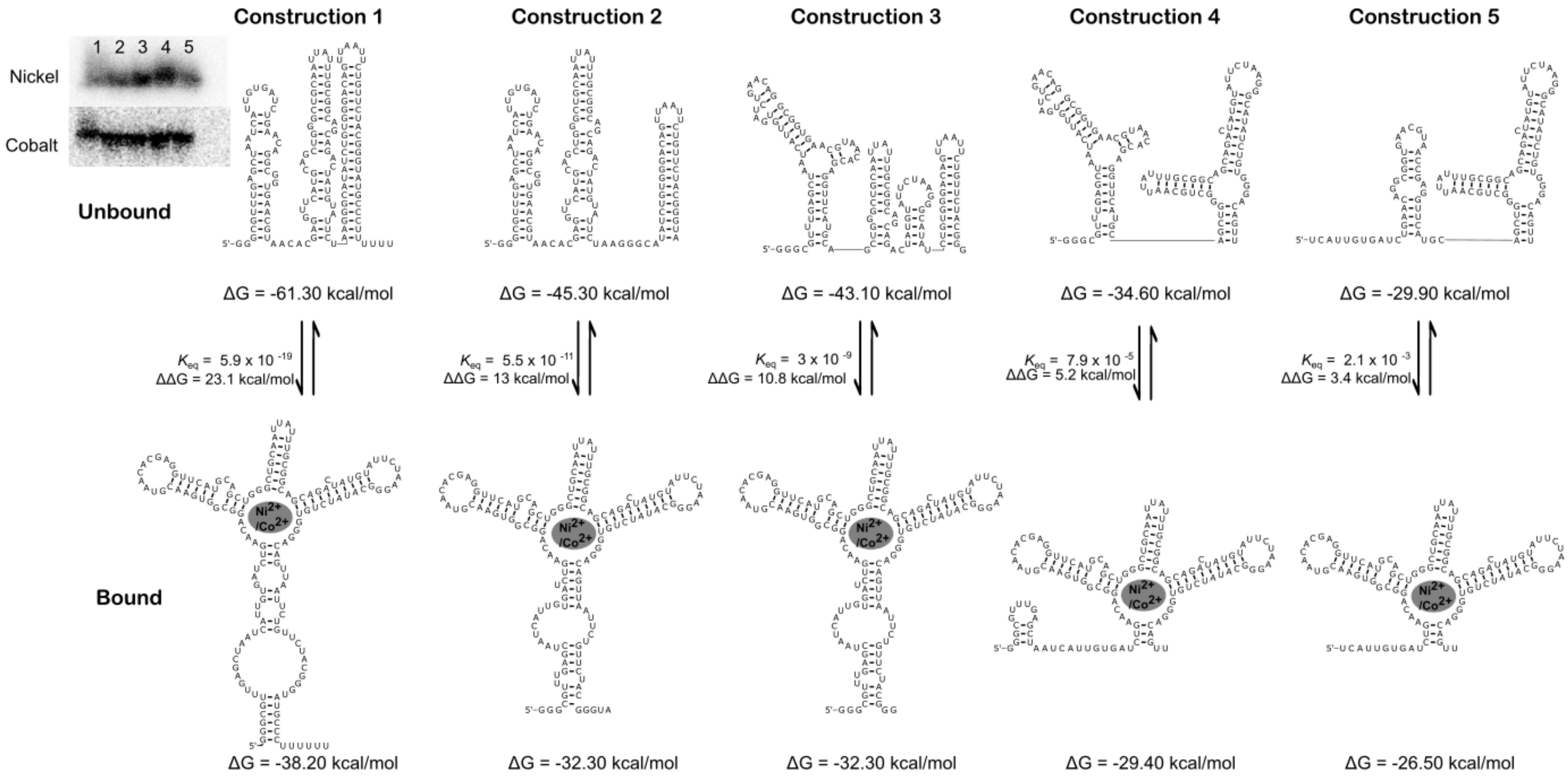

**Supplementary Fig. 5 | Free energy of all different constructions of the nickel-cobalt riboswitch used in the SR-PAGE experiment in their bound (constrained) and unbound (unconstrained) conformations.**

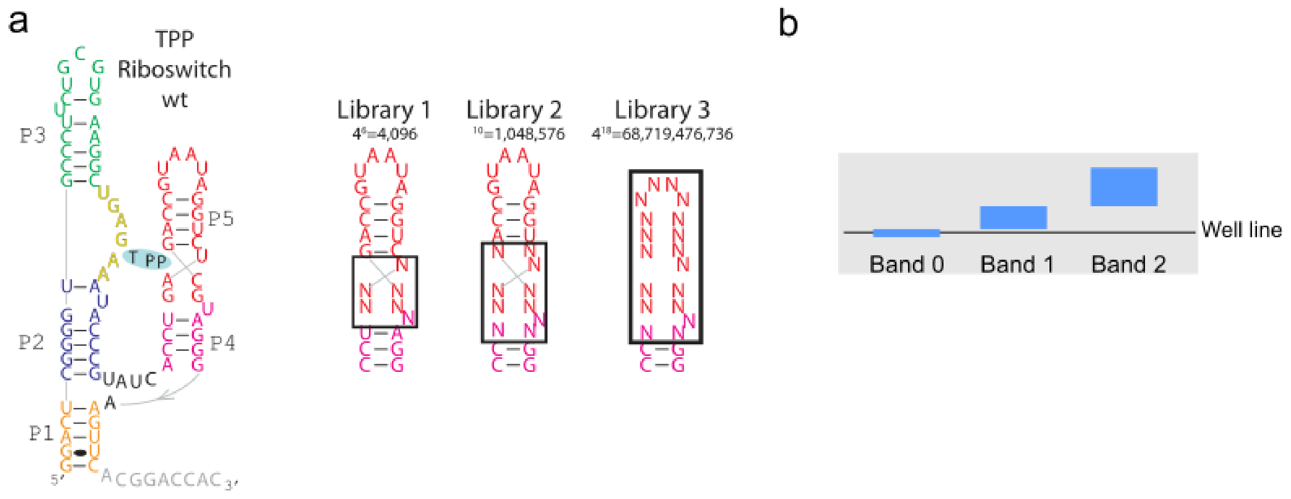

**Supplementary Fig. 6 | Degenerated libraries of TPP riboswitch**

**a** Libraries 1, 2 and 3 had 6, 10 and 18 degenerated nucleotides respectively within stem P5. **b** Schematic representation of the bands that were cut-out after the second migration of the SR-PAGE (Supplementary Fig.8c). The cut 0 is only the wells, The cut 1 is approximately from 2 mm to 1 cm above the wells and the cut 2 is approximately 1 cm above the wells and above

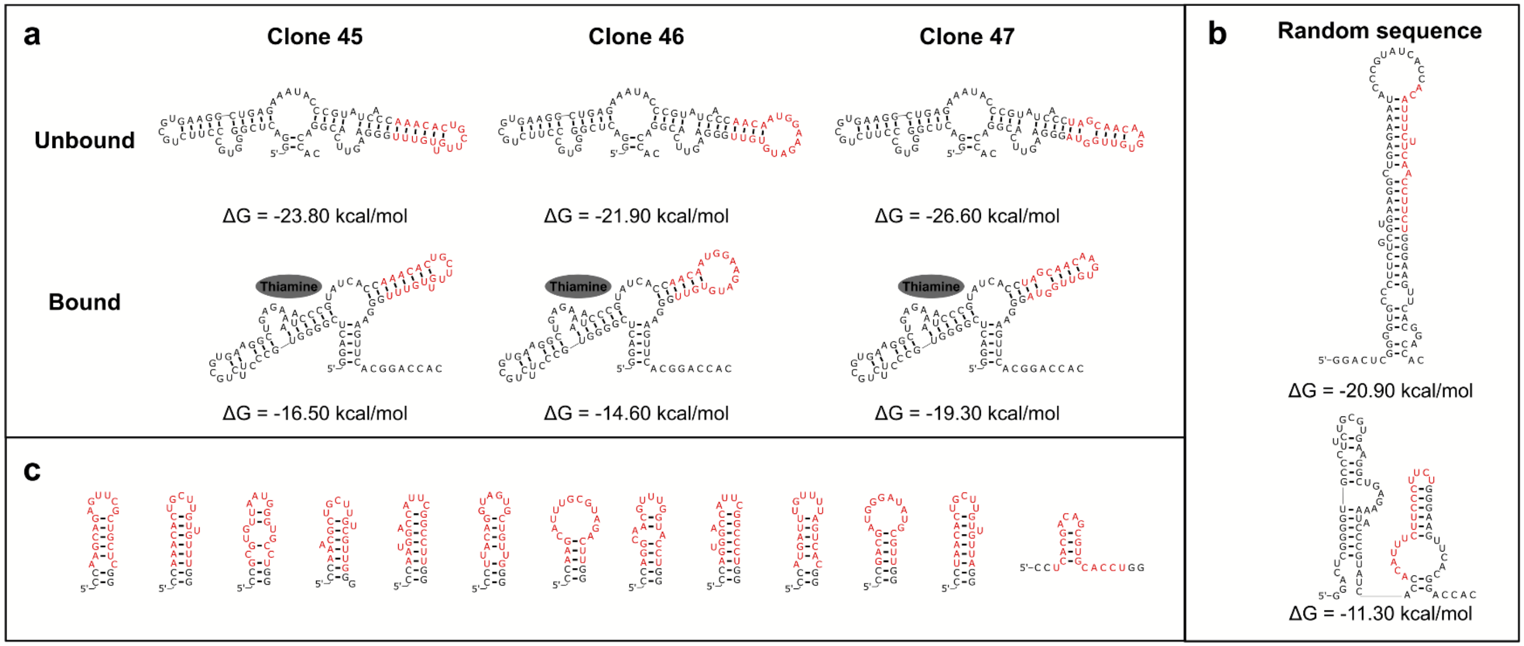

**Supplementary Fig. 7 | Library 3 TPP-derived thiamine switches have a stem that replaces P4-P5.**

**a**, Secondary structures were prepared as for Supplementary Figs. 1-4 by constraining only P2 and P3, between which the thiamine-binding pocket is found. It is noteworthy that the three validated sequences derived from the selected library 3 have similar overall bound/unbound structures, i.e. where the randomized regions (in red) can form a stem present in both the predicted bound and unbound states. **b**, Secondary structure of a randomly generated sequence expected to be found in the starting pool, with the same constraints as in **a**. **c**,The representativity of these examples was further evaluated by looking at the sequences with four or more reads in our deep sequencing. The vast majority of selected sequences can form stems. Conversely, only a minority of non-selected random sequences can form stems (Supplementary Table 4).

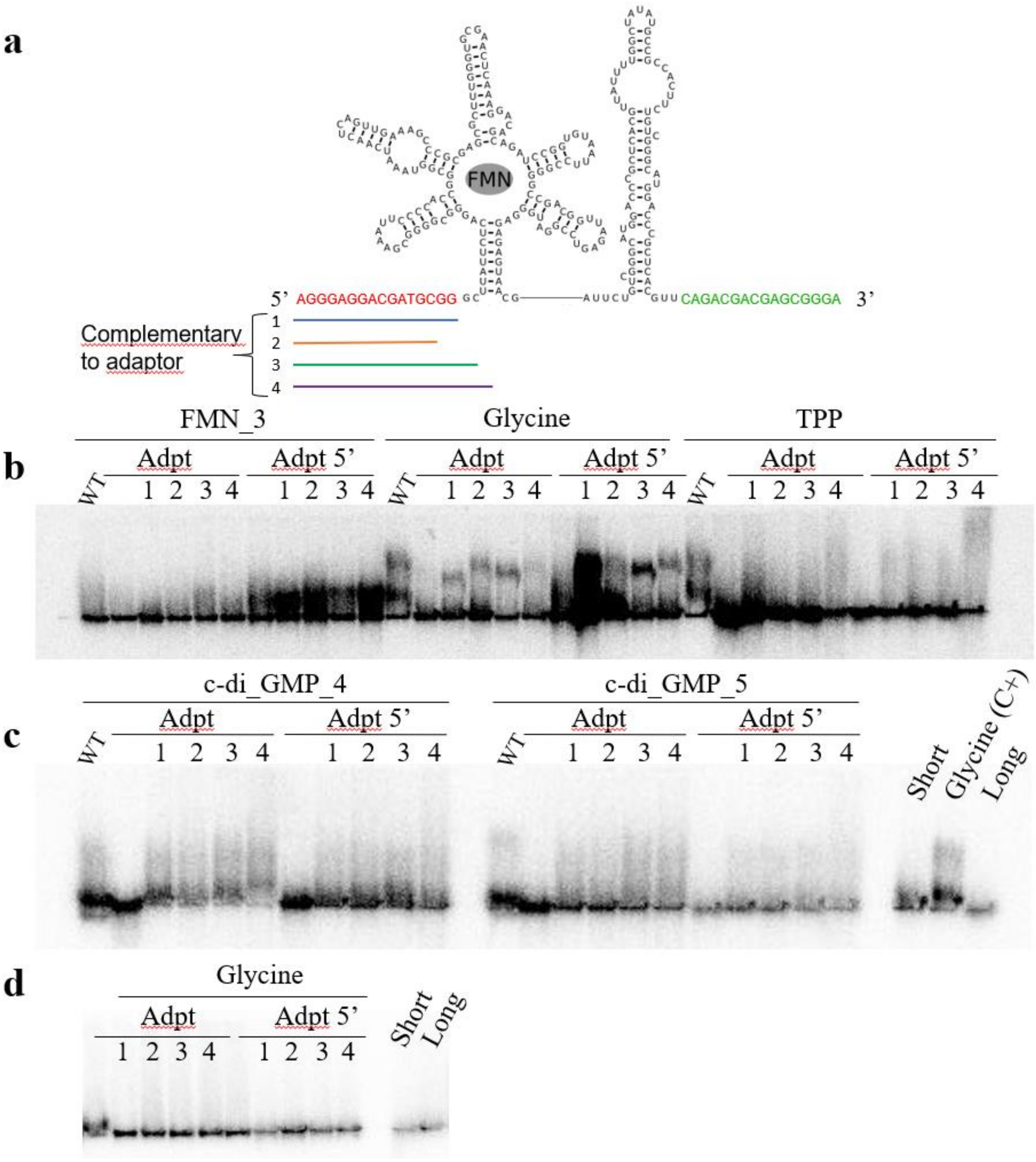

**Supplementary Fig. 8 | The presence of oligonucleotides complementary to the adapters makes it possible to restore the shift of the riboswitches by the SR-PAGE method.**

**a**, Schematic of the design of complementary oligonucleotides with an example on the FMN riboswitch. The sequences in red and green are the adapters for PCR amplification. The colored lines represent the different complementary oligonucleotides tested: 1 is fully complementary to the adapter; 2 is complementary to the adapter minus two nucleotides; 3 is fully complementary to the adapter with two additional nucleotides complementary to the riboswitch sequence; and 4 is fully complementary to the adapter with four additional nucleotides complementary to the riboswitch sequence. **b**, Result of SR-PAGE with FMN_3, Glycine and TPP riboswitches by spraying the corresponding ligand. “Adpt’’ corresponds to the 5' and 3' complementary adapters. “Adpt 5’’ corresponds to the 5' complementary adapter only. “WT’’ is the riboswitch without adapters. The numbers correspond to the combination of the oligonucleotides as described before. **c**, Result of SR-PAGE with c-di-GMP_4 and _5 with a solution of c-di-GMP sprayed. ‘’Short’’, ‘’Long ‘’ represents two controls to verify proper gel migration. Glycine riboswitch is used as a positive control, so the ligand glycine was also sprayed. **d**, Result of SR-PAGE with glycine riboswitch without spraying any ligand.

### Supplementary Tables

#### Supplementary Table 1 | List of all the oligonucleotides.

| **Degenerated libraries of the TPP riboswitch** |
|---|
| **Librairie 1 _SELEX_TPP**<br>- Lib1_For: TAATACGACTCACTATAggactcggggtgcccttctgcgtgaaggctgagaaatacccgtatcacc<br>- Lib1_Rev: gtggtccgtgaacttccctNNNctggcattatccagNNNaggtgatacgggtatttctc |
| **Librairie 2_SELEX_TPP**<br>- Lib2_For: TAATACGACTCACTATAggactcggggtgcccttctgcgtgaaggctgagaaatacccgtatcacc<br>- Lib2_Rev: gtggtccgtgaacttcccNNNNNtggcattatccaNNNNNggtgatacgggtatttctc |
| **Librairie 3_SELEX_TPP**<br>- Lib3_For: TAATACGACTCACTATAggactcggggtgcccttctgcgtgaaggctgagaaatacccgtatcacc<br>- Lib3_Rev: gtggtccgtgaacttcccnnnnnnnnnnnnnnnnnggtgatacgggtatttctc |
| **Amplification from clones pGEMT (amplified from miniprep)**<br>- Cloning_For: TAATACGACTCACTATAggactcggggtgccctt<br>- Cloning_Rev: gtggtccgtgaacttccc<br>(note: full sequence of these clones available in Supplementary Table 2) |
| **Enriched clones (Assembly PCR)**<br>- TPP_For: TAATACGACTCACTATAggactcggggtgcccttctgcgtgaaggctgagaaatacccgtatcacc<br>- TPP_Rev:<br> ○ TPP1_Rev: gtggtccgtgaacttccctaggctggcattatccaggcgaggtgatacgggtatttctc<br> ○ TPP2_Rev: gtggtccgtgaacttcccttgtctggcattaccaggtcaggtgatacgggtatttctc<br> ○ TPP3_Rev: gtggtccgtgaacttcccgagcatggcattatccaaacaaggtgatacgggtatttctc<br> ○ TPP4_Rev: gtggtccgtgaacttcccaagcatggcattatccaaacaaggtgatacgggtatttctc<br> ○ TPP5_Rev: gtggtccgtgaacttcccagcattggcattatccaaaaccggtgatacgggtatttctc<br> ○ TPP6_Rev: gtggtccgtgaacttcccaaccatggcattatccaaagcaggtgatacgggtatttctc<br> ○ TPP7_Rev: gtggtccgtgaacttcccttcgctggcattatccagacgaggtgatacgggtatttctc<br> ○ TPP8_Rev: gtggtccgtgaacttccctgctctggcattatccaactggggtgatacgggtatttctc<br> ○ TPP45_Rev: gtggtccgtgaacttcccaaacaacaagcagtgtttggtgatacgggtatttctc<br> ○ TPP46_Rev: gtggtccgtgaacttcccaacacatcttccattgttggtgatacgggtatttctc |

- TPP47_Rev: gtggtccgtgaacttccctaccaacacttgttgctaggtgatacgggtatttctc
- TPP48_Rev: gtggtccgtgaacttcccgagcgtccgatctgcttcggtgatacgggtatttctc

**Fluorescent oligonucleotide (control for the SR-PAGE migration)**

Cy3-tccccttgcatacatgtcgcgtttatgcttcgaccctctagccgcaccccttgg as well as

Cy3-ccatcggcactgacggcctcacttgcagcagaacgtgggcagcttgctgaatcgttctgcaagagtgagcccgtaacataatgg

**Primers for constructions of FMN riboswitch (Amplified from *Escherichia coli*)**

**FMN_For**: TAATACGACTCACTATAGGgcttattctcagggcgg

**FMN_1**

- FMN_1_Rev: attatggttaccagaatcagggca
- FMN_1_full sequence:
  TAATACGACTCACTATAGGgcttattctcagggcggggcgaaattccccaccggcggtaaatcaactcagttgaaagcccgcgAgcgctttgggtgcgaactcaaaggacagcagatccggtgtaattccggggccgacggttagagtccggatgggagagagtaacgattctgtcgggcatggacccgctcacgttattttggctatatgccgccactcctaagactgccctgattctggtaaccataat

**FMN_2**

- FMN_2_Rev: cagggcagtcttaggagt
- FMN_2_full sequence:
  TAATACGACTCACTATAGGgcttattctcagggcggggcgaaattccccaccggcggtaaatcaactcagttgaaagcccgcgagcgctttgggtgcgaactcaaaggacagcagatccggtgtaattccggggccgacggttagagtccggatgggagagagtaacgattctgtcgggcatggacccgctcacgttattttggctatatgccgccactcctaagactgccctg

**FMN_3**

- FMN_3_Rev: gtggcggcatatagccaa
- FMN_3_full sequence:
  TAATACGACTCACTATAGGgcttattctcagggcggggcgaaattccccaccggcggtaaatcaactcagttgaaagcccgcgagcgctttgggtgcgaactcaaaggacagcagatccggtgtaattccggggccgacggttagagtccggatgggagagagtaacgattctgtcgggcatggacccgctcacgttattttggctatatgccgccac

**FMN_4**

- FMN_4_Rev: aacgtgagcgggtccat
- FMN_4_full sequence:
  TAATACGACTCACTATAGGgcttattctcagggcggggcgaaattccccaccggcggtaaatcaactcagttgaaagcccgcgagcgctttgggtgcgaactcaaaggacagcagatccggtgtaattccggggccgacggttagagtccggatgggagagagtaacgattct

gtcgggcatggacccgctcacgtt

**FMN_5**

- FMN_5_Rev: tcgttactctctcccatcc
- FMN_5_full sequence:

TAATACGACTCACTATAGGgcttattctcagggcggggcgaaattccccaccggcggtaaatcaactcagttgaaagcccgcg
agcgctttgggtgcgaactcaaaggacagcagatccggtgtaattccggggccgacggttagagtccggatgggagagagtaacga

**Primers for constructions of Fluoride riboswitch (Amplified from *Burkholderia thailandensis*)**

**Fluor_For**: TAATACGACTCACTATAGGgcggctaccggagatgg

**Fluor_1**

- Fluor_1_Rev: atggcctacgacctcctg
- Fluor_1_full sequence:

TAATACGACTCACTATAGGgcggctaccggagatggcatgcctccgtacaaccgccggcgagccggctgatgatgcctacgc
gttcctgggtgcaggaggtcgtaggccat

**Fluor_2**

- Fluor_2_Rev: tacgacctcctgcaccc
- Fluor_2_full sequence:

TAATACGACTCACTATAGGgcggctaccggagatggcatgcctccgtacaaccgccggcgagccggctgatgatgcctacgc
gttcctgggtgcaggaggtcgta

**Fluor_3**

- Fluor_3_Rev: cctcctgcacccagg
- Fluor_3_full sequence:

TAATACGACTCACTATAGGgcggctaccggagatggcatgcctccgtacaaccgccggcgagccggctgatgatgcctacgc
gttcctgggtgcaggagg

**Fluor_4**

- Fluor_4_Rev: tgcacccaggaacgc
- Fluor_4_full sequence:

TAATACGACTCACTATAGGgcggctaccggagatggcatgcctccgtacaaccgccggcgagccggctgatgatgcctacgc
gttcctgggtgca

| |
|---|
| **Fluor_5**<br>- Fluor_5_Rev: aggaacgcgtaggcatca<br>- Fluor_5_full sequence:<br>TAATACGACTCACTATAGGgcggctaccggagatggcatgcctccgtacaaccgccggcgagccggctgatgatgcctacgc gttcct |
| **Primers for construction of c-di-GMP I riboswitch (PCR assembly of riboswitch from *Vibrio cholerae*)** |
| **c-di-GMP_1** (created by PCR assembly and used as template for other GMP constructions)<br>- c-di-GMP_1_For_A: TAATACGACTCACTATAGGgaaaaatgtcacgcacagggc<br>- c-di-GMP_1_Rev_B: ggtttaggccggaggctttgcgtcccactctttcgaatggtttgccctgtgcgtg<br>- c-di-GMP_1_For_C: ctccggcctaaaccagaagacatggtaggtagcggggttaccgatggcaaaatgcataca<br>- c-di-GMP_1_Rev_D: aaagcatgcattcatagtgtcaatgatgagtcaacaaagtgtatgcattttgccatcg<br>- c-di-GMP_1_full sequence:<br>TAATACGACTCACTATAGGggaaaaatgtcacgcacagggcaaaccattcgaaagagtgggacgcaaagcctccggccta aaccagaagacatggtaggtagcggggttaccgatggcaaaatgcatacactttgttgactcatcattgacactatgaatgcatgcttt |
| **c-di-GMP_2**<br>- c-di-GMP_for: TAATACGACTCACTATAGGgaaaaatgtcacgcacagggc<br>- c-di-GMP_2_Rev: gtgtcaatgatgagtcaacaaagt<br>- c-di-GMP_2_full sequence:<br>TAATACGACTCACTATAGGggaaaaatgtcacgcacagggcaaaccattcgaaagagtgggacgcaaagcctccggccta aaccagaagacatggtaggtagcggggttaccgatggcaaaatgcatacactttgttgactcatcattgacac |
| **c-di-GMP_3**<br>- c-di-GMP_3_Rev: tgatgagtcaacaaagtgtatgc<br>- c-di-GMP_3_full sequence:<br>- TAATACGACTCACTATAGGggaaaaatgtcacgcacagggcaaaccattcgaaagagtgggacgcaaagcctccggccta aaccagaagacatggtaggtagcggggttaccgatggcaaaatgcatacactttgttgactcatca |
| **c-di-GMP_4**<br>- c-di-GMP_4_Rev: aaagtgtatgcattttgccatc<br>- c-di-GMP_4_full sequence:<br>TAATACGACTCACTATAGGggaaaaatgtcacgcacagggcaaaccattcgaaagagtgggacgcaaagcctccggccta |

aaccagaagacatggtaggtagcggggttaccgatggcaaaatgcatacactttg

**c-di-GMP_5**

- c-di-GMP_5_Rev: gtatgcattttgccatcggta
- c-di-GMP5_full sequence:
- TAATACGACTCACTATAGGggaaaaatgtcacgcacagggcaaaccattcgaaagagtgggacgcaaagcctccggcctaaaccagaagacatggtaggtagcggggttaccgatggcaaaatgcatac

**Primers for construction of glycine riboswitch (PCR assembly of riboswitch from *Vibrio cholerae*)**

- Gly_For_A: TAATACGACTATAGGgttgaagactgcaggagagtggttgttaaccagattttaacatctgagccaaataacccg
- Gly_Rev_B: tcgcttattcgttgccaatatatggctaagaataatgcacctgaaagatttacttcttcggcgggttatttggctcagatg
- Gly_For_C: ttggcaacgaataagcgaggactgtagttggaggaacctctggagagaaccgtttaatcggtcgccgaaggagcaag
- Gly_Rev_D: tcctctgtccttttgcctgagagtttcactctgcatatgcgcagagcttgctccttcggcgaccgat
- Gly_full sequence:

  TAATACGACTATAGGgttgaagactgcaggagagtggttgttaaccagattttaacatctgagccaaataacccgccgaagaagtaAatctttcaggtgcattattcttagccatatattggcaacgaataagcgaggactgtagttggaggaacctctggagagaaccgtttaatcggtcgccgaaggagcaagctctgcgcatatgcagagtgaaactctcaggcaaaaggacagagga

**Primers for construction of TPP riboswitch (Assembly PCR of TPP riboswitch from *Escherichia coli*)**

- TPP_For: TAATACGACTCACTATAggactcggggtgcccttctgcgtgaaggctgagaaatacccgtatcacc
- TPP_Rev: gtggtccgtgaacttccctacggctggcattatccagatcaggtgatacgggtatttctc
- TPP_full sequence:

  TAATACGACTCACTATAGGggactcggggtgcccttctgcgtgaaggctgagaaatacccgtatcacctgatctggataatgccagcgtagggaagttcacggaccac

**Primers for construction of nickel-cobalt riboswitch (PCR assembly of riboswitch from *Listeria monocytogenes*)**

**NiCo_1 (template NiCo_4)**

- NiCo_1_F: TTCTAATACGACTCACTATAGGgcgtttgagctaatcattgtgatctgaacaggcg
- NiCo_1_R: aaaaaagggcatacccgtagaacagaattaac
- NiCo_1_full_sequence:

  TTCTAATACGACTCACTATAGGgcgtttgagctaatcattgtgatctgaacaggcggtgaacgtaacacgaggttcatgcagctgggctgcaattatttgcggcagcagactatgtattctaagggcatatctgtgggacagttaattctgttctacgggtatgcccttttt

**NiCo_2 (template NiCo_4)**

- NiCo_2_F: TTCTAATACGACTCACTATAGGgcgtttgagctaatcattgtgatctgaacaggcg
- NiCo_2_R: tacccgtagaacagaattaactgtcccacagatatgc
- NiCo_2_full_sequence:
  TTCTAATACGACTCACTATAGGgcgtttgagctaatcattgtgatctgaacaggcggtgaacgtaacacgaggttcatgcagct
  gggctgcaattatttgcggcagcagactatgtattctaagggcatatctgtgggacagttaattctgttctacgggta

**NiCo_3 (template NiCo_4)**

- NiCo_3_F: TTCTAATACGACTCACTATAGGgcgtttgagctaatcattgtgatctgaacaggcg
- NiCo_3_R: cccgtagaacagaattaactgtcccacagatatgc
- NiCo_3_full_sequence:
  TTCTAATACGACTCACTATAGGgcgtttgagctaatcattgtgatctgaacaggcggtgaacgtaacacgaggttcatgcagct
  gggctgcaattatttgcggcagcagactatgtattctaagggcatatctgtgggacagttaattctgttctacggg

**NiCo_4 (template Nico_5)**

- NiCo_4_F: TTCTAATACGACTCACTATAGGgcgtttgagctaatcattgtgatctgaacaggcg
- NiCo_4_R: aactgtcccacagatatgccctt
- NiCo_4_full_sequence:
  TTCTAATACGACTCACTATAGGgcgtttgagctaatcattgtgatctgaacaggcggtgaacgtaacacgaggttcatgcagct
  gggctgcaattatttgcggcagcagactatgtattctaagggcatatctgtgggacagtt

**NiCo_5 (created by PCR assembly and used as template for other nickel-cobalt constructions)**

- NiCo_1_For_A: TTCTAATACGACTCACTATAGGtcattgtgatctgaacaggcggtgaacgtaa
- NiCo_1_Rev_B: tgcagcccagctgcatgaacctcgtgttacgttcaccgcctg
- NiCo_1_For_C: cagctgggctgcaattatttgcggcagcagactatgtattctaagggcatatctgtggga
- NiCo_1_Rev_D: aactgtcccacagatatgccctt
- NiCo_1_full_sequence:
  TTCTAATACGACTCACTATAGGtcattgtgatctgaacaggcggtgaacgtaacacgaggttcatgcagctgggctgcaattat
  ttgcggcagcagactatgtattctaagggcatatctgtgggacagtt

**Test of the influence of adapters**

**c-di-GMP adapter (to synthesize the riboswitch with few combinations of adapters)**

**Template control riboswitch:**

- c-di-GMP_For_noT7: ggaaaaatgtcacgcacaggg
- c-di-GMP4_Rev: caaagtgtatgcattttgccatc
- c-di-GMP5_Rev: gtatgcattttgccatcggta

**c-di-GMP4_adpt 5'_3'**:

- c-di-GMP4_adapt 5'_3'_For: gaaaTTAATACGACTCACTATAGGGA*gggaggacgatgcgg*ggaaaaatgtcacgcacaggg
- c-di-GMP4_adapt 5'_3'_Rev : *tcccgctcgtcgtctgc*aaagtgtatgcattttgccatc

**c-di-GMP4_adpt 5'**:

- c-di-GMP4_adapt 5'_For:

  gaaaTTAATACGACTCACTATAGGGA*gggaggacgatgcgg*ggaaaaatgtcacgcacaggg
- c-di-GMP4_adapt 5'_Rev : caaagtgtatgcattttgccatc

**c-di-GMP4_adpt 3'**:

- c-di-GMP4_adapt 3'_For:

  TTAATACGACTCACTATAgggaaaaatgtcacgcacagggc
- c-di-GMP4_adapt 3'_Rev : *tcccgctcgtcgtctgc*aaagtgtatgcattttgccatc

**c-di-GMP5_adpt 5'_3'**:

- c-di-GMP5_adapt 5'_3'_For : gaaaTTAATACGACTCACTATAGGGA*gggaggacgatgcgg*ggaaaaatgtcacgcacaggg
- c-di-GMP5_adapt 5'_3'_Rev : *tcccgctcgtcgtctg*gtatgcattttgccatcggta

**c-di-GMP5_adpt 5'**:

- c-di-GMP5_adapt 5'_For:

  gaaaTTAATACGACTCACTATAGGGA*gggaggacgatgcgg*ggaaaaatgtcacgcacaggg
- c-di-GMP4_adapt 5'_Rev : gtatgcattttgccatcggta

**c-di-GMP5_adpt 3'**:

- c-di-GMP5_adapt 3'_For: TTAATACGACTCACTATAgggaaaaatgtcacgcacagggc
- c-di-GMP5_adapt 3'_Rev : *tcccgctcgtcgtctgc*gtatgcattttgccatcggta

**Glycine adapter (to synthesize the riboswitch with few combinations of adapters)**

**Template:**

- Gly_For_noT7: gttgaagactgcaggagag
- Gly_Rev: tcctctgtccttttgcctga

**Gly_adpt 5'_3’**:

- Gly_adapt 5’_3’_For: gaaaTTAATACGACTCACTATAGGGA*gggaggacgatgcgg*gttgaagactgcaggagag
- Gly_adapt 5’_3’_Rev : *tcccgctcgtcgtctg*tcctctgtccttttgcctga

**Gly_adpt 5'**:

- Gly_adapt 5’_For: gaaaTTAATACGACTCACTATAGGGA*gggaggacgatgcgg*gttgaagactgcaggagag
- Gly_adapt 5’_Rev : tcctctgtccttttgcctga

**Gly_adpt 3'**:

- Gly_adapt 3’_For: TTAATACGACTCACTATAgg
- Gly_adapt 3’_Rev : *tcccgctcgtcgtctg*tcctctgtccttttgcctga

**FMN adapter (to synthesize the riboswitch with few combinations of adapters)**

**Template**:

- FMN_For_noT7: gcttattctcagggcgg
- FMN_Rev: gtggcggcatatagccaa

**FMN_adpt 5'_3’**:

- FMN_adapt 5’_3’_For: gaaaTTAATACGACTCACTATAGGGA*gggaggacgatgcgg*gcttattctcagggcgg
- FMN_adapt 5’_3’_Rev : *tcccgctcgtcgtctg*gtggcggcatatagccaa

**FMN_adpt 5’**:

- FMN_adapt 5’_For: gaaaTTAATACGACTCACTATAGGGA*gggaggacgatgcgg*gcttattctcagggcgg
- FMN_adapt 5’_Rev : gtggcggcatatagccaa

**FMN_adpt 3’**:

- FMN_adapt 3’_For: TAATACGACTCACTATAgggcttattctcagggcgg
- FMN_adapt 5’_Rev : *tcccgctcgtcgtct*ggtggcggcatatagccaa

**TPP adapter (to synthesize the riboswitch with few combinations of adapters)**

**Template**:

- TPP_For_noT7: ggactcggggtgccctt
- TPP_Rev: gtggtccgtcaagtccct

**TPP_adpt 5'_3'**:

- TPP_adapt 5'_3'_For: gaaaTTAATACGACTCACTATAGGGA*gggaggacgatgcgg*ggactcggggtgccctt
- TPP_adapt 5'_3'_Rev : *tcccgctcgtcgtctg*gtggtccgtcaagttccct

**TPP_adpt 5'**:

- TPP_adapt 5'_For: gaaaTTAATACGACTCACTATAGGGA*gggaggacgatgcgg*ggactcggggtgccctt
- TPP_adapt 5'_Rev : gtggtccgtcaagtccct

**TPP_adpt 3'**:

- TPP_adapt 3'_For: TTAATACGACTCACTATAGgg
- TPP_adapt 3'_Rev : *tcccgctcgtcgtctg*gtggtccgtcaagttccct

**Primers complementary to adapter sequences for all tested riboswitches**

- all_adpt_5' (1): ccgcatcgtcctccc
- all_adpt_3' (1): tcccgctcgtcgtctg
- Adpt_2_5' (2): gcatcgtcctccc
- Adpt-2_3' (2): tcccgctcgtcgtc

**Primers complementary to adapter sequences for c-di-GMP riboswitch**

- Adpt_c-di-GMP 4/5 + 2nts_5'(3): ccccgcatcgtcctccc
- Adpt_c-di-GMP 4/5 + 4nts_5'(3): ttccccgcatcgtcctccc
- Adpt_c-di-GMP4 + 4nts_3' (3 & 4): tcccgctcgtcgtctgcaaa
- Adpt_c-di-GMP5 + 4nts_3' (3 & 4): tcccgctcgtcgtctggtat

**Primers complementary to adapter sequences for glycine riboswitch**

- Adpt_Gly + 2nts_5' (3): acccgcatcgtcctccc
- Adpt_Gly + 4nts_5'(4): caacccgcatcgtcctccc
- Adpt_Gly + 4nts_3' (3 & 4): tcccgctcgtcgtctgtcct

**Primers complementary to adapter sequences for FMN riboswitch**

- Adpt_FMN+ 2nts_5' (3): gcccgcatcgtcctccc
- Adpt_FMN + 4nts_5'(4): aagcccgcatcgtcctccc
- Adpt_FMN + 4nts_3' (3 & 4): tcccgctcgtcgtctggtgg

| Primers complementary to adapter sequences for glycine riboswitch | |
|---|---|
| - | Adpt_TPP + 2nts_5' (3): cccgcatcgtcctccc |
| - | Adpt_TPP + 4nts_5' (4): gtccccgcatcgtcctccc |
| - | Adpt_TPP + 4nts_3' (3 & 4): tcccgctcgtcgtctggtgg |

The capital letters correspond to the sequence of the T7 promoter. The letter ‘’N’’ represents degenerated nucleotides. The abbreviation ‘’For’’ corresponds to forward primers, whereas ‘’Rev’’ stands for reverse primers. The underlined nucleotides correspond to the aptamer of each riboswitch. The italicized nucleotides correspond to the adapters, either in 5’ or 3’ end of the riboswitch

**Supplementary Table 2 | List of constraints applied to Mfold software**

| Riboswitch | Constraints |
|---|---|
| Fluoride | ............xx(((((..........xxxx(((((....)))))xxxxx)))))xx......... |
| FMN | ..((((((((..((.((((.......)))).)).((((....(((((...)))))....))))..((.((((((((....))))))))....))...(((((...... .))))) .(((.(((.......)))..)))..))))))))... |
| c-di_GMP I | ..........((((xxxxxx((...((((((....))))))...))xxx(((.(((((((...(((.......))).))))..))))))xxx)).)).... ......... |
| Nickel-cobalt | ..........(((.....((...(((............))).....))...((.((........)).))..((((...................)))).....))).. |
| Stem P2-P3, Library 3 TPP | .....((((((((((((.....)))))))......))))))).......................................... |

For the realization of Fig. 3 and Supplementary Figs. 2-5 and 7. Dots represent nucleotides with no constraints. "X" shows nucleotides that are forced to remain unbound, whereas parentheses indicate base pairs.

**Supplementary Table 3 | Clones selected with the SELEX of the degenerated TPP riboswitch.**

<table>
<tr><th>Clone from</th><th>Clone number</th><th>Lib.(see Fig. 5a)</th><th>Select in band (see Fig. 5b)</th><th>$K_D$ Thiamine (µM)</th><th>$K_D$ TPP (µM)</th><th>Ratio $K_D^T$/ $K_D^{TPP}$</th></tr>
<tr><td rowspan="16">Illumina sequencing</td><td>1</td><td>1</td><td>2</td><td>17.7</td><td>67.1</td><td>0.26</td></tr>
<tr><td colspan="6">RNA sequence:<br>GGACUCGGGGUGCCCUUCUGCGUGAAGGCUGAGAAAUACCCGUAUCACCU<br>CGCCUGGAUAAUGCCAGCCUAGGGAAGUUCACGGACCAC</td></tr>
<tr><td>2</td><td>1</td><td>2</td><td>16.5</td><td>18.5</td><td>0.88</td></tr>
<tr><td colspan="6">RNA sequence:<br>GGACUCGGGGUGCCCUUCUGCGUGAAGGCUGAGAAAUACCCGUAUCACCU<br>GACCUGGAUAAUGCCAGACAAGGGAAGUUCACGGACCAC</td></tr>
<tr><td>3</td><td>2</td><td>2</td><td>/</td><td>/</td><td>/</td></tr>
<tr><td colspan="6">RNA sequence:<br>GGACUCGGGGUGCCCUUCUGCGUGAAGGCUGAGAAAUACCCGUAUCACCUU<br>GUUUGGAUAAUGCCAUGCUCGGGAAGUUCACGGACCAC</td></tr>
<tr><td>4</td><td>2</td><td>2</td><td>22.9</td><td>21</td><td>1.09</td></tr>
<tr><td colspan="6">RNA sequence:<br>GGACUCGGGGUGCCCUUCUGCGUGAAGGCUGAGAAAUACCCGUAUCACCU<br>UGUUUGGAUAAUGCCAUGCUUGGGAAGUUCACGGACCAC</td></tr>
<tr><td>5</td><td>2</td><td>2</td><td>/</td><td>/</td><td>/</td></tr>
<tr><td colspan="6">RNA sequence:<br>GGACUCGGGGUGCCCUUCUGCGUGAAGGCUGAGAAAUACCCGUAUCACCG<br>GUUUUGGAUAAUGCCAAUGCUGGGAAGUUCACGGACCAC</td></tr>
<tr><td>6</td><td>2</td><td>2</td><td>/</td><td>/</td><td>/</td></tr>
<tr><td colspan="6">RNA sequence:<br>GGACUCGGGGUGCCCUUCUGCGUGAAGGCUGAGAAAUACCCGUAUCACCU<br>GCUUUGGAUAAUGCCAUGGUUGGGAAGUUCACGGACCAC</td></tr>
<tr><td>7</td><td>2</td><td>1</td><td>/</td><td>/</td><td>/</td></tr>
<tr><td colspan="6">RNA sequence:<br>GGACUCGGGGUGCCCUUCUGCGUGAAGGCUGAGAAAUACCCGUAUCACCU<br>CGUCUGGAUAAUGCCAGCGAAGGGAAGUUCACGGACCAC</td></tr>
<tr><td>8</td><td>2</td><td>1</td><td>/</td><td>/</td><td>/</td></tr>
<tr><td colspan="6">RNA sequence:<br>GGACUCGGGGUGCCCUUCUGCGUGAAGGCUGAGAAAUACCCGUAUCACCC<br>GAGUUGGAUAAUGCCAGAGCAGGGAAGUUCACGGACCAC</td></tr>
<tr><td></td><td>9</td><td>1</td><td>0</td><td>34.12</td><td>34.55</td><td>0.99</td></tr>
</table>

<table>
<tr><td rowspan="20">Generation 4</td><td colspan="6">RNA sequence:<br>GGACUCGGGGUGCCCUUCUGCUGAAGGCUGAGAAAUACCCGUAUCACCU<br>GUACUGGAUAUGCCAGAAUAGGGAAGUUCACGGACCAC</td></tr>
<tr><td>10</td><td>1</td><td>0</td><td>/</td><td>/</td><td>/</td></tr>
<tr><td colspan="6">RNA sequence :<br>GGACUCGGGGUGCCCUUCUGCGUGAAGGCUGAGAAAUACCCGUAUCACCU<br>ACACUGGAAAUGCCAGACAAGGGAAGUUCACGGACCAC</td></tr>
<tr><td>11</td><td>1</td><td>0</td><td>/</td><td>/</td><td>/</td></tr>
<tr><td colspan="6">RNA sequence :<br>GGACUCGGGGUGCCCUUCUGCGUGAAGGCUGAGAAAUACCCGUAUCACCU<br>GUUCUGGAUAAUGCAGUAUAGGGAAGUUCACGGACCAC</td></tr>
<tr><td>12</td><td>1</td><td>1</td><td>/</td><td>/</td><td>/</td></tr>
<tr><td colspan="6">RNA sequence :<br>GGACUCGGGGUGCCCUUCUGCGUGAAGGCUGAGAAAUACCCGUAUCACCU<br>AGUCUGGAAAUGCCAGAUGAGGGAAGUUCACGGACCAC</td></tr>
<tr><td>13</td><td>1</td><td>1</td><td>2.59</td><td>9.27</td><td>0.28</td></tr>
<tr><td colspan="6">RNA sequence:<br>GGACUCGGGGUGCCCUUCUGCGUGAAGGCUGAGAAAUACCCGUAUCACCU<br>GUACUGGAUAAUGCCAGCGAGGGAAGUUCACGGACCAC</td></tr>
<tr><td>14</td><td>1</td><td>1</td><td>14.86</td><td>13.46</td><td>1.1</td></tr>
<tr><td colspan="6">RNA sequence:<br>GGACUCGGGGUGCCCUUCUGCGUGAAGGCUGAGAAAUACCCGUAUCACCU<br>CUGCUGGAUAAUGCCAGGAUAGGGAAGUUCACGGACCAC</td></tr>
<tr><td>15</td><td>1</td><td>2</td><td>/</td><td>/</td><td>/</td></tr>
<tr><td colspan="6">RNA sequence:<br>GGACUCGGGGUGCCCUUCUGCGUGAAGGCUGAGAAAUACCCGUAUACCU<br>GUGCUGGUAAUGCCAGUUCAGGGAAGUUCACGGACCAC</td></tr>
<tr><td>16</td><td>1</td><td>2</td><td>/</td><td>/</td><td>/</td></tr>
<tr><td colspan="6">RNA sequence:<br>GGACUCGGGGUGCCCUUCUGCGUGAAGGCUGAGAAAUACCCGUAUCACCU<br>AUUCUGGAUAAUGCCAGUAAGGGAAGUUCACGGACCAC</td></tr>
<tr><td>17</td><td>1</td><td>2</td><td>/</td><td>/</td><td>/</td></tr>
<tr><td colspan="6">RNA sequence:<br>GGACUCGGGGUGCCCUUCUGCGUGAAGGCUGAGAAAUACCCGUAUCACCU<br>GACUGGUAAUGCCAGUAUAGGGAAGUUCACGGACCAC</td></tr>
<tr><td>18</td><td>2</td><td>0</td><td>58</td><td>50.4</td><td>1.15</td></tr>
<tr><td colspan="6">RNA sequence:<br>GGACUCGGGGUGCCCUUCUGCGUGAAGGCUGAGAAAUACCCGUAUCACCU<br>CGCUUGGAUAUGCCACAUGAGGGAAGUUCACGGACCAC</td></tr>
<tr><td>19</td><td>2</td><td>0</td><td>45</td><td>33.1</td><td>1.36</td></tr>
</table>

| | | | | | | |
|---|---|---|---|---|---|---|
| | RNA sequence: GGACUCGGGGUGCCCUUCUGCGUGAAGGCUGAGAAAUACCCGUAUCACCU CGCUUGGAUAUGCCACAUGAGGGAAGUUCACGGACCAC | | | | | |
| | 20 | 2 | 0 | / | / | / |
| | RNA sequence: GGACUCGGGGUGCCCUUCUGCGUGAAGGCUGAGAAAUACCCGUAUCACCU GGAUUGGAUAAUGCCAAGUUGGGAAGUUCACGGACCAC | | | | | |
| | 21 | 2 | 1 | / | / | / |
| | RNA sequence: GGACUCGGGGUGCCCUUCUGCGUGAAGGCUGAGAAAUACCCGUAUCACCU GUGCUGGAUAAUGCCAGAGUAGGGAAGUUCACGGACCAC | | | | | |
| | 22 | 2 | 1 | / | / | / |
| | RNA sequence: GGACUCGGGGUGCCCUUCUGCGUGAAGCUGAGAAAUACCCGUAUCACCCAU GAUGGAUAAUGCCAAUAAGGGGAAGUUCACGGACCAC | | | | | |
| | 23 | 2 | 1 | 12.5 | 40.2 | 0.31 |
| | RNA sequence: GGACUCGGGGUGCCCUUCUGCGUGAAGGCUGAGAAAUACCCGUAUCACCUA AAUUGGAUAAUGCCACUAGAGGGAAGUUCACGGACCAC | | | | | |
| | 24 | 2 | 2 | 17.53 | 20.11 | 0.87 |
| | RNA sequence: GGACUCGGGGUGCCCUUCUGCGUGAAGGCUGAGAAAUACCCGUAUCACCU AUAUUGGAUAAUGCCACCUAAGGGAAGUUCACGGACCAC | | | | | |
| | 25 | 2 | 2 | 3.07 | 5.1 | 0.6 |
| | RNA sequence: GGACUCGGGGUGCCCUUCUGCGUGAAGGCUGAGAAAUACCCGUAUCACC AAGGAUGGAUAAUGCCACUAAGGGGAAGUUCACGGACCAC | | | | | |
| | 26 | 2 | 2 | / | / | / |
| | RNA sequence: GGACUCGGGGUGCCCUUCUGCGUGAAGGCUGAGAAAUACCCGUAUCACC UAAACUGGAUAAUGCCAUAAAGGGGAAGUUCACGGACCAC | | | | | |
| | 27 | 1 | 0 | / | / | / |
| | RNA sequence: GGACUCGGGGUGCCCUUCUCGUGAAAGAAGAGAAAUCCCGUAUCACCUU UGCUGGAAAUGUCAGCUUAGGGAAGUUCACGGACCAC | | | | | |
| | 28 | 1 | 0 | / | / | / |
| | RNA sequence: GGACUCGGGGUGCCCUUCUGCCGAAGGCUGAGAAAUACCCGUAUCACCUA GACUGGAUAAUCCAGCCCAGGGAAGUUCACGGACCAC | | | | | |
| | 29 | 1 | 0 | / | / | / |

| | | | | | | |
|---|---|---|---|---|---|---|
| Generation 10 | RNA sequence: GGACUCGGGGUGCCCUUCUGUGCUGAGAAAGGAAUACCCGUAUACCUAU GCUGGAAAUGCCAGUGCAGGGAAGUUCACGGAC | | | | | |
| | 30 | 1 | 1 | / | / | / |
| | RNA sequence: GGACUCGGGGUGCCCUUCUGCUGAAGGCUGAGAAAUACCCGUAUCACCU UGUCUGUAAUGCAGUCGAGGGAAGUUCACGGACCAC | | | | | |
| | 31 | 1 | 1 | / | / | / |
| | RNA sequence: GGCUCGGGGUGCCCUUCUCGUGAAGCUGAGAAAUACCCGUAUCACCUAAA CUGGUAAUGCCAGUCUAGGGAAGUUCACGGACCAC | | | | | |
| | 32 | 1 | 1 | / | / | / |
| | RNA sequence: GGACUCGGGGUGCCCUUCUGGUGAAGGUGAGAAAUCCUGUAUCACCUGAUC UGGAUAAUGCCGCCCAGGGAAGUUCACGGACCAC | | | | | |
| | 33 | 1 | 2 | / | / | / |
| | RNA sequence: GGACUCGGGGUGCCCUUCUGCGUGAAGGCUGAGAAAUACCCGUACACCUAC ACUGGAUAAUGCCAGCUAAGGGAAGUUCACGGACCAC | | | | | |
| | 34 | 1 | 2 | 69 | 39 | 1.77 |
| | RNA sequence: GGACUCGGGGUGCCCUUCUGCGUGAAGGCUGAGAAAUACCCGUAUCACCU GAAGAUGUUUCUGCUCGGGGAAGUUCACGGACCAC | | | | | |
| | 35 | 1 | 2 | / | / | / |
| | RNA sequence: GGACUCGGGGUGCCCUUCUGGUGAAGGUGAGAAAUCCUGUAUCACCUGAUC UGGAUAAUGCCGCCCAGGGAAGUUCACGGACCAC | | | | | |
| | 36 | 2 | 0 | / | / | / |
| | RNA sequence: GGACUCGGGGUGUCUUCCUCGUGAGGCUGAGAAUAUCCGAUACCAGACUCC | | | | | |
| | 37 | 2 | 0 | / | / | / |
| | RNA sequence: GGACUCGGGGUGGCGNCCUGCGUGAAGCUGAGAAUCUCGUUCUCUAAAGCC CN | | | | | |
| | 38 | 2 | 0 | / | / | / |
| | RNA sequence: GGACUCGGGGUGUCCUUCUGCGUGAGGCUGAGAAUCNUCGUUCACUCCCG ACCCNN | | | | | |
| | 39 | 2 | 1 | / | / | / |
| | RNA sequence: GGACUCGGGGUGCCCUUCUGCGUGAAGGCUGAGAAAUACCCGUAUCACCUG | | | | | |

| | | | | | | |
|---|---|---|---|---|---|---|
| | ACUUGGAUAAUGCCACUUGGGGGAAGUUCACGGACCAC | | | | | |
| | 40 | 2 | 1 | / | / | / |
| | RNA sequence:<br>GGACUCGGGGUGCCCUUCUGCGUGAAGGCUGAGAAAUACCCGCAUCACCAGCACUGGUAAUGCCAAUACUGGGAAGUUCACGGACCAC | | | | | |
| | 41 | 2 | 1 | 71 | 168 | 0.42 |
| | RNA sequence:<br>GGACUCGGGGUGCCCUUCUGCGUGAAGGCUGAGAAAUACCCGUAUUACCGCGUGUGGAUAAUCCAGCGUCGGGAAGUUCAC | | | | | |
| | 42 | 2 | 2 | 60 | 140 | 0.43 |
| | RNA sequence<br>:GGACUCGGGGUGCCCUUCUGCGUGAAGGCUGAGAAAUACCCGUAUCACCAACGAACGCUAGAACGUUGGGAAGUUCACGGACCAC | | | | | |
| | 43 | 2 | 2 | 6.1 | 6.7 | 0.91 |
| | RNA sequence:<br>GGACUCGGGGUGCCCUUCUGCGUGAAGGCUGAGAAAUACCCGUAUCACCACGAAUGGAUAAUGCCAGGCUGGGAAGUUCACGGACCAC | | | | | |
| | 44 | 2 | 2 | 8.9 | 25 | 0.36 |
| | RNA sequence:<br>GGACUCGGGGGUGCCCUUCUGCGUGAAGGCUGAGAAAUACCCGUAUCACCCAAACUGGAUAAUGCCAUCCGGGGGAAGUUCACGGACCACAAUCAC | | | | | |
| Illumina sequencing | 45 | 3 | 2 | 3.7 | 1000* | 0.004 |
| | RNA sequence:<br>GGACUCGGGGUGCCCUUCUGCGUGAAGGCUGAGAAAUACCCGUAUCACCAAACACUGCUUGUUGUUUGGGAAGUUCACGGACCAC | | | | | |
| | 46 | 3 | 2 | 77 | 262 | 0.29 |
| | RNA sequence:<br>GGACUCGGGGUGCCCUUCUGCGUGAAGGCUGAGAAAUACCCGUAUCACCAACAAUGGAAGAUGUGUUGGGAAGUUCACGGACCAC | | | | | |
| | 47 | 3 | 2 | 5.89 | 1000* | 0.005 |
| | RNA sequence:<br>GGACUCGGGGUGCCCUUCUGCGUGAAGGCUGAGAAAUACCCGUAUCACCUAGCAACAAGUGUUGGUAGGGAAGUUCACGGACCAC | | | | | |

Many sequences were assayed by in-line probing without providing conclusive results, in such cases, the $K_D$ is indicated as “/”, this is presumably due to lack of binding to the ligand or a modulation too weak to be quantified above background, as estimated by in-line probing. Also, we have assayed sequences from the initial libraries (prior to selection), but none showed modulation. For the affinity with TPP of clones 45 and 47 (represented with an asterisk, we could not measure the $K_{D,}$ since no modulation was observed. We put a value of 1 mM, which was the largest tested concentration.

**Supplementary Table 4 | Predicted stem formation in the random region of library 3.**

| Seq ID | 5′-3′ | dG value | GC% (within 18 bp) | Final GC pairing occurs or not (position) |
|---|---|---|---|---|
| >22_1 | CCAAGCAGAGTTCGCTGCTCGG | -7.80 | 56% | Yes (1,22) |
| >6_2 | CCAAACACTGCTTGTTGTTTGG | -6.30 | 33% | Yes (1,22) |
| >6_3 | CCGCGTGTTAATGGGTGCCTGG | -2.60 | 56% | Yes (1,22) |
| >5_5 | CCAAACGCTGCTTGTCGTTGGG | -4.40 | 50% | Yes (1,21) |
| >5_6 | CCAATGGACCATTCGGCCTTGG | -6.70 | 50% | Yes (1,22) |
| >5_4 | CCTTACAGGTAGTGCTGTTGGG | -6.10 | 44% | Yes (1,22) |
| >4_11 | CCAAGCATTTGCGTAGACTTGG | -2.80 | 39% | Yes (1,22) |
| >4_9 | CCAGGCAACGTTTGGTACCTGG | -6.10 | 50% | Yes (1,22) |
| >4_7 | CCAGTGGACCATTCGGCCCTGG | -9.10 | 61% | Yes (1,22) |
| >4_8 | CCATGATTTGTTTTAGTCACGG | -4.10 | 28% | Yes (1,22) |
| >4_10 | CCGACGATGGGATATGCGTTGG | -5.40 | 50% | Yes (1,22) |
| >4_13 | CCTAACACTGCTTGTTGTTAGG | -6.70 | 33% | Yes (1,22) |
| >4_12 | CCTCACGACAGCGTGCACCTGG | -2.90 | 61% | No |
| >3_17 | CCTATGATTGCTCAGTCGTAGG | -9.50 | 39% | Yes (1,22) |
| >3_22 | CCTATGCATCCGAGACACAAGG | 0.30 | 44% | Yes (1,22) |
| >3_14 | CCTCCATCAGACGGGATGGCGG | -8.20 | 61% | Yes (1,22) |
| >3_23 | CCTCTGATGAGTGTCAGAAGGG | -8.40 | 44% | No |
| >3_28 | CCTGATAGACTAGTGCTAAGGG | -1.70 | 39% | No |
| >3_18 | CCTGATTAGCACTGCGAAATGG | 0.30 | 39% | Yes (1,22) |
| >3_33 | CCTTATGTTGGAGTAGCTAGGG | -3.50 | 39% | Yes (1,22) |
| >3_29 | CCTTTTGTTGCGCAGTTTTAGG | 0.00 | 33% | Yes (1,22) |
| >3_69 | CCAACAATGGAAGATGTGTTGG | -4.40 | 33% | Yes (1,22) |
| >3_71 | CCAACACGTGTCTGATTGTTGG | -4.30 | 39% | Yes (1,22) |
| >3_73 | CCAACAGTGTCTCGCTGCTTGG | -7.10 | 50% | Yes (1,22) |
| >3_75 | CCAACGAATTGATGTCGGTTGG | -3.90 | 39% | Yes (1,22) |
| >3_77 | CCAAGATTCGCACCGATCTTGG | -7.90 | 44% | Yes (1,22) |
| >3_79 | CCAATCATGGAAGATGGGTTGG | -4.10 | 39% | Yes (1,22) |
| >3_81 | CCAATGCTCTTGACGGTGTTGG | -6.70 | 44% | Yes (1,22) |
| >3_83 | CCACCAGTTTACGGCTGGCGGG | -10.00 | 61% | No |
| >3_85 | CCACCTCATGGGTATGAGGGGG | -9.70 | 56% | Yes (1,22) |
| >3_87 | CCACGAAGAACAGTGTCGAGGG | -2.90 | 50% | No |
| >3_89 | CCACGCACTGTTGGGGTGCTGG | -8.20 | 61% | Yes (1,22) |
| >3_91 | CCACGGAATGGCTAACCGCAGG | -1.60 | 56% | Yes (1,22) |
| >3_93 | CCACTCATGGCTCATGTGTTGG | -3.70 | 44% | Yes (1,22) |
| >3_95 | CCACTGATTCTTGTTGAAAGGG | -1.10 | 53% | No |
| >3_97 | CCAGACAGGATGCTGTCATGGG | -9.60 | 61% | No |
| >3_99 | CCAGACTGCTGTTGCCGCCTGG | -5.30 | 70% | Yes (1,22) |
| >3_101 | CCAGATCTACCGTTGGATTTGG | -8.50 | 57% | Yes (1,22) |
| >3_103 | CCAGCCCTACTTTGTCATATGG | 0.50 | 56% | Yes (1,22) |
| >3_105 | CCAGCGGCTGCTGCATGCCTGG | -6.60 | 74% | Yes (1,22) |
| >3_107 | CCAGGCCACTTCTGTTACCTGG | -7.50 | 50% | Yes (1,22) |
| >3_109 | CCAGGCTGGAGCCCTTGCTTGG | -8.40 | 61% | Yes (1,22) |
| >3_111 | CCAGGTGCGGTCTAAAACCAGG | -2.60 | 50% | Yes (1,22) |
| >3_113 | CCCAAATGCCAATTGGCCAGGG | -6.50 | 50% | Yes (1,22) |
| >3_115 | CCCCATGCGTTACGTTTGGAGG | -5.70 | 50% | Yes (1,22) |
| >3_117 | CCCCTACAAGAGACTGCTGGGG | -6.80 | 56% | Yes (1,22) |
| >3_119 | CCCTACCATCTGCTTGCGAGGG | -4.70 | 56% | Yes (1,22) |

| | | | | |
|---|---|---|---|---|
| >3_121 | CCCTACCTGAGCTAGTTAAGGG | -2.60 | 44% | Yes (1,22) |
| >3_123 | CCCTTGCCGCTGGTATGTCGGG | -3.70 | 61% | Yes (1,22) |
| >3_125 | CCCTTGGACACTTTCCGGGGGG | -12.20 | 61% | No |
| >3_127 | CCGAAGGCTTGTGCTTAGTAGG | -3.20 | 44% | Yes (1,22) |
| >3_129 | CCGCTTGATCCGTATATGCCGG | -0.90 | 50% | No |
| >3_131 | CCGGACAGCACTCGCTACCCGG | -8.60 | 67% | Yes (1,22) |
| >3_133 | CCGGAGCATTCCTGTGCCCCGG | -10.80 | 67% | Yes (1,22) |
| >3_135 | CCGGCTGTTCTGTACCGCCCGG | -5.60 | 67% | Yes (1,22) |
| >3_137 | CCGGGATCGCTCGACGCCCCGG | -8.30 | 78% | Yes (1,22) |
| >3_139 | CCGTCAGCCACGTCGCTTAAGG | -2.70 | 56% | Yes (1,22) |
| >3_141 | CCGTTGATGCTTTCGCAACCGG | -3.30 | 50% | Yes (1,22) |
| >3_143 | CCTAAGCCCATCCAACCTTAGG | -3.90 | 44% | Yes (1,22) |
| >3_145 | CCTAAGCGTTGTCGCGCCTGGG | -8.10 | 61% | Yes (1,22) |
| >3_147 | CCTAGATGCTCTTAGCATGGGG | -5.80 | 44% | Yes (1,22) |
| >3_149 | CCTAGCGGTTTGCAAAGTCGGG | -2.60 | 50% | Yes (1,22) |
| >3_151 | CCTAGTTCTGGATCGCACTGGG | -4.30 | 50% | Yes (1,22) |
| >3_153 | CCTATCGCTGAGTGCGCCTGGG | -5.60 | 61% | Yes (1,22) |
| >3_155 | CCTCCACGCTCGGATTACCCGG | -3.00 | 61% | No |
| >3_157 | CCTCTTATTTCACCGAAGGAGG | -4.50 | 39% | Yes (1,22) |
| >3_159 | CCTGAACCAGACGCGAATTTGG | -2.50 | 44% | No |
| >3_161 | CCTGAAGGTAGGAATACTAGGG | -3.60 | 39% | No |
| >3_163 | CCTGAATGCACGTAGCTCATGG | -2.10 | 44% | Yes (1,22) |
| >3_165 | CCTGACCATCTGGGTTGTTGGG | -5.40 | 50% | Yes (1,22) |
| >3_167 | CCTGACTTGCGTGTACCTTGGG | -1.50 | 50% | Yes (1,22) |
| >3_169 | CCTGAGATTTGCCGTCTCGAGG | -6.50 | 50% | Yes (1,22) |
| >3_171 | CCTGCGAATGCGCGTTCATGGG | -4.90 | 56% | Yes (1,22) |
| >3_173 | CCTGCTCGTTGGTAGTGCCAGG | -3.10 | 56% | No |
| >3_175 | CCTGGACGTTTTACACGCTGGG | -6.20 | 50% | Yes (1,22) |
| >3_177 | CCTGGTGTTTGGCTATGCTGGG | -6.50 | 50% | Yes (1,22) |
| >3_179 | CCTGTCGATGTTCGTCGATGGG | -10.70 | 50% | Yes (1,22) |
| >3_181 | CCTGTTAGATGTGCTCTACAGG | -6.40 | 39% | Yes (1,22) |
| >3_183 | CCTGTTCACGGGCGTGCGTAGG | -6.60 | 61% | Yes (1,22) |
| >3_185 | CCTTCACTATTGCTGGGATGGG | -3.20 | 44% | No |
| >3_187 | CCTTCCACACTGCCGCACCTGG | 0.50 | 61% | No |
| >3_189 | CCTTCCGGAGAGGAGCCATCGG | -3.00 | 61% | No |
| >3_191 | CCTTGCACAATGCTGTGCGGGG | -12.00 | 56% | Yes (1,22) |
| >3_193 | CCTTGTCATCTGCTGTGCTGGG | -1.80 | 50% | Yes (1,22) |
| >3_195 | CCTTGTTGCAATTGGCGCAGGG | -7.40 | 50% | Yes (1,22) |
| >3_197 | CCTTTGCGCGTTGGGTGCCTGG | -5.00 | 61% | Yes (1,22) |
| Random Seq No | Random Seq (5'-3') | dG | GC% (within 18 bp) | Final GC pairing occurs or not (position) |
| 1. | ccTTCCCTTGCATATATGTTgg | -0.90 | 33% | No |
| 2. | ccACATTTCTTCAACCTTCTgg | -0.20 | 33% | No |
| 3. | ccAATTGCACCCTTAGGACGgg | -2.60 | 50% | No |
| 4. | ccAAGACAGATATGTTCTTAgg | -3.00 | 28% | Yes (1, 22) |
| 5. | ccCCTATATTTCATCATTGGgg | -5.00 | 33% | Yes (1,22) |
| 6. | ccCAACGGGATCGCATGTCCgg | -5.90 | 61% | No |
| 7. | ccCACGTAAAACATTGTTAAgg | 1.00 | 28% | No |
| 8. | ccACCCTCAGGTTTTTGAGCgg | -4.90 | 50% | No |
| 9. | ccGACAAAAACTTTAAAAAGgg | 0.30 | 22% | No |
| 10. | ccAAATTCGCGCTCATAACTgg | 1.10 | 39% | Yes (1,22) |
| 11. | ccGTTAGGCCACGATTGCGTgg | -5.90 | 56% | No |
| 12. | ccGAGTTTCGGCCCTGTGCTgg | -3.50 | 61% | No |

| | | | | |
|---|---|---|---|---|
| 13. | ccGCGCTGTATAGCCGATTCgg | -3.40 | 56% | No |
| 14. | ccTCATTCGGGCCTTATATCgg | -0.60 | 44% | No |
| 15. | ccTGGAAACCCCAACCTATTgg | -1.70 | 44% | No |
| 16. | ccTAGACAGCATCATTGGCCgg | 0.10 | 50% | No |
| 17. | ccGAAGTTATTGGGCATATTgg | -2.00 | 33% | No |
| 18. | ccCACCGTAAAGTCCTCCTCgg | -1.00 | 56% | No |
| 19. | ccGGGCGTCCCTCCTTTAAAgg | -3.30 | 56% | No |
| 20. | ccAGATGATAAGCTCCGGCAgg | -1.50 | 50% | No |
| 21. | ccAAGGATCGGTGATATTAAgg | 0.50 | 33% | No |
| 22. | ccCAAAGATTCGGCACATTAgg | 0.10 | 39% | No |
| 23. | ccCTCTTGTTGGTGTGGTATgg | -0.30 | 44% | No |
| 24. | ccCGCTTAACTGCGTGGCGGgg | -10.20 | 67% | No |
| 25. | ccAGCCTTATGGCAAAATCGgg | -5.00 | 44% | No |
| 26. | ccTTCGGGAATGATTCTGGTgg | -5.80 | 44% | Yes (1,22) |
| 27. | ccAACGCTAAAGGTCCATAGgg | 0.40 | 44% | No |
| 28. | ccCACATACATCGCAACCTGgg | -2.00 | 50% | Yes (1,22) |
| 29. | ccGCATGCGTTCAATTTGACgg | -2.00 | 44% | No |
| 30. | ccGATCGCTTGGCGCTAAGAgg | -1.10 | 56% | No |
| 31. | ccTTAAAGCGGCTGCACTGCgg | -4.30 | 56% | No |
| 32. | ccTGTAAGGACGATTACGGAgg | -5.00 | 44% | No |
| 33. | ccGTGGGCGGCCTGGGGGGAgg | -5.00 | 83% | No |
| 34. | ccGCACTACCCCATCGACCTgg | -0.60 | 61% | No |
| 35. | ccGTACAGGAACACTCTATAgg | -1.00 | 39% | No |
| 36. | ccTTGCTCTCAGACGAACAAgg | -5.70 | 44% | Yes (1,22) |
| 37. | ccATTACTAGAGTGCCGCTTgg | -0.50 | 44% | No |
| 38. | ccTCAGCCCCCCTGTCGTCGgg | -3.00 | 72% | No |
| 39. | ccCGTTGTTGTGATTCGACTgg | -3.70 | 44% | Yes (1,22) |
| 40. | ccCTATTGAGGCATCAACTGgg | -6.60 | 44% | Yes (1,22) |
| 41. | ccAATGAATCGGCCTATGTCgg | -3.00 | 44% | No |
| 42. | ccCCCGATGTCGTTAGTGAAgg | 0.20 | 50% | No |
| 43. | ccGGTTCCGACGCATACCTCgg | -2.80 | 61% | No |
| 44. | ccCTTCGTTGAGAACCCACAgg | 0.00 | 50% | No |
| 45. | ccATCATACAACTGGGGACAgg | -5.80 | 44% | No |
| 46. | ccTAATCCCTACGCCCATCAgg | 0.40 | 50% | No |
| 47. | ccTCTACACGCGTCTCTGTGgg | -3.40 | 56% | No |
| 48. | ccGCTCCAGTTCATGTGCTGgg | -6.80 | 56% | No |
| 49. | ccGGAGAGCACCCTCCACAAgg | -5.90 | 61% | No |
| 50. | ccGGTCTAGTGGTATGGTGGgg | -3.10 | 56% | No |
| 51. | ccTGATACACGCGGCAGGGGgg | -3.60 | 67% | No |
| 52. | ccTAGGACCATCGGTAGTAGgg | -6.80 | 50% | No |
| 53. | ccCTGACCACTGCCTATAGGgg | -4.20 | 56% | No |
| 54. | ccAGAGTGTCAGCCAGTGTAgg | 0.40 | 50% | No |
| 55. | ccACCCCACGAGGATCCGAGgg | -5.00 | 67% | No |
| 56. | ccAAGGCGAACCGGGCCAGAgg | -4.30 | 67% | No |
| 57. | ccCTCAAAGCCGCGGCGCGAgg | -7.60 | 72% | No |
| 58. | ccAGTAGCCCCGGGGTGAACgg | -4.70 | 67% | Yes (1,22) |
| 59. | ccACCTATGGGGCTGGATAAgg | -4.40 | 50% | No |
| 60. | ccAACTGCCCTGGTGAGCGCgg | -4.10 | 67% | No |
| 61. | ccTCTGCTGCTCGAGGCCGTgg | -4.00 | 67% | No |
| 62. | ccTCGCCGATGCTTGCTGCGgg | -3.90 | 67% | No |
| 63. | ccTCCCCAGCCGCTACATCTgg | -1.90 | 61% | No |
| 64. | ccGTCTCTTTGCCGACTAATgg | -1.90 | 44% | No |
| 65. | ccGCGAACAACCACACCATAgg | 0.30 | 50% | No |

| | | | | |
|---|---|---|---|---|
| 66. | ccGCGATTCGTCGGGGCGCCgg | -5.20 | 78% | No |
| 67. | ccTCGGAATACGGTATGGGCgg | -1.60 | 56% | No |
| 68. | ccTCGCGGACGCCAGGCATCgg | -4.50 | 72% | No |
| 69. | ccGTGCAGGTAGCGGAGGCCgg | -4.20 | 72% | No |
| 70. | ccCGCACGCGAGACGAACTGgg | -2.90 | 67% | Yes (1,22) |

We observed that selected sequences could form stems within the randomized region. To evaluate whether it actually occurred much more often than it would for random sequences, we evaluated the average delta-G of structures for that portion of sequence for random sequences (as would be found in the initial library, prior to selection with SR-PAGE) vs our selected sequences. Random sequences (18nt in size, with additional cc in 5' and gg in 3') were created using the "Random DNA Sequence Generator" http://www.faculty.ucr.edu/~mmaduro/random.htm with an input of seven different values of GC percentage in the sequences (0.35, 0.40, 0.45, 0.5, 0.55, 0.60 and 0.65) and to evaluate its folding nature through structure formation, whether dG value and terminal GC base pairs occurred or not. Folding was performed with Mfold.

It is noteworthy that some of our sequenced samples selected from library 3 were actually contaminations from library 2 (sequences which started dominating the population of library 2 earlier, given its smaller complexity than library 3). These sequences were excluded from the above analysis (given their different size and larger number of fixed nucleotides, they were easily recognizable, a total of 625,731 reads corresponding to the correct size for library 3 were obtained from selected pools), but the fact that we selected them as contaminants from library 2 actually supports our interpretation that using SR-PAGE, we selected for the presence of a stem at a position roughly equivalent to P4 and P5.
